# Multicellular dynamics on structured surfaces: Stress concentration is a key to controlling complex microtissue morphology on engineered scaffolds

**DOI:** 10.1101/2023.03.06.530721

**Authors:** Ryosuke Matsuzawa, Akira Matsuo, Shuya Fukamachi, Sho Shimada, Midori Takeuchi, Takuya Nishina, Philip Kollmannsberger, Ryo Sudo, Satoru Okuda, Tadahiro Yamashita

## Abstract

Tissue engineers have utilized a variety of three-dimensional (3D) scaffolds for controlling multicellular dynamics and the resulting tissue microstructures. In particular, cutting-edge microfabrication technologies, such as 3D bioprinting, provide increasingly complex structures. However, unpredictable microtissue detachment from scaffolds, which ruins desired tissue structures, is becoming an evident problem. To overcome this issue, we elucidated the mechanism underlying collective cellular detachment by combining a new computational simulation method with quantitative tissue-culture experiments. We first quantified the stochastic processes of cellular detachment shown by vascular smooth muscle cells on model curved scaffolds and found that microtissue morphologies vary drastically depending on cell contractility, substrate curvature, and cell-substrate adhesion strength. To explore this mechanism, we developed a new particle-based model that explicitly describes stochastic processes of multicellular dynamics, such as adhesion, rupture, and large deformation of microtissues on structured surfaces. Computational simulations using the developed model successfully reproduced characteristic detachment processes observed in experiments. Crucially, simulations revealed that cellular contractility-induced stress is locally concentrated at the cell-substrate interface, subsequently inducing a catastrophic process of collective cellular detachment, which can be suppressed by modulating cell contractility, substrate curvature, and cell-substrate adhesion. These results show that the developed computational method is useful for predicting engineered tissue dynamics as a platform for prediction-guided scaffold design.

## 1. Introduction

Various microfabrication technologies have been employed in tissue engineering to create scaffolds with precise structures at micrometre and even nanometre scales [1–4]. Such scaffolds are designed and utilised to serve as a structural template that guides cells to autonomously form the types of hierarchical structures found *in vivo* [5–7]. Recent studies in bioengineering and mechanobiology have clarified that intrinsic cellular force and its interaction with geometrical cues from the surrounding environment have a crucial impact on cellular behaviour [8–13]; the influence of such forces is especially manifested in mechanical events such as collective cellular migration [14–16] and alignment [17–19]. These studies highlight the importance of a rational geometrical design of cell-culture scaffolds that correctly direct cellular behaviour based upon quantitative prediction, to construct a tissue with the desired structure and biological properties *in vitro*. There are, however, so far only limited means of quantitatively describing how cell-generated force and geometrical constraints determine the fate of tissues growing in artificial environments.

In the last decade, curved surfaces have drawn considerable attention as model cell-culture environments in this field as they offer the chance to explore how geometrical constraints influence cellular behaviours in three dimensions [20–25]. In particular, the effect of surface curvature on the growth kinetics of microtissues confined in microspaces has been extensively investigated in the context of bone tissue formation [26,27] and wound healing [28,29]. Dunlop *et al.* pointed out that there are two distinct stages in such engineered tissue formation processes [30]. First, cells form a confluent monolayer on the scaffold surface, where the chemical composition and microscopic topographies such as the roughness of the substrate surface have a major influence on cellular behaviours. Second, the confluent cell sheet grows to form a thick and matured microtissue on the scaffold, where the macroscopic substrate geometry, including curvature at supracellular scale, characterises the growth kinetics. Experimental studies have elucidated the basic tendency for the speed of the microtissue growth in the second stage to be proportional to the local curvature of the growing front [31]; this tendency has since been explained by several different theoretical models [32,33]. The basic concept of curvature-dependent microtissue growth was applied to predict tissue formation processes in through holes of various geometries [34] in a model 3D environment. Technical improvements including level-set method for detecting the local curvature in three dimensions [35] have further widened the potential for simulating microtissue growth in 3D environments of increasing complexity such as beam-based lattices [36]; these methods are likely to serve as a theoretical basis to optimise the scaffold geometry for practical use in the growing field of 3D bioprinting [37]. It is notable that tissue formation process *in vivo* also follows the rules deduced by these theoretical models, which thus seem to offer considerable potential for computation-aided scaffold design in tissue engineering in the future [38]. Nevertheless, these models are built on the strong assumption that the growing tissue firmly attaches to the scaffold surface and never causes ruptures; this is not always the case.

Cells in a growing microtissue exert a considerable amount of contractile force on the surrounding extracellular matrix (ECM) [39–41]. This allows cells to alter the surrounding microenvironment to regulate the proper growth steps including ECM deposition and phenotype control [42–44]. Several experimental studies have demonstrated that this kind of contractile cellular behaviour often drives unpredictable deformation of tissues growing on structured surfaces, including rupture, aggregation and detachment from the scaffold surface [45–48]. Such phenomena undermine the basic premises of conventional models that attempt to predict microtissue growth kinetics on microstructures. Thus, a quantitative framework predicting the stability of cell-substrate interface, *i.e.*, whether contractile cells will succeed in forming a stable layer on structured surfaces or cause a critical transition from confluent cell layer to 3D tissue is desirable as a fundamental basis for modelling an entire microtissue formation process *in silico*. To date, no modelling discipline has been established that can describe such large deformations of the viscoelastic microtissue body together with dynamic rearrangements of cell-cell and cell-substrate adhesions. Furthermore, the fundamental mechanism by which cellular force drives such multicellular dynamics under complex geometrical constraints remains unclear. The problem in understanding and predicting microtissue deformation is the lack of an experimental system that enables quantitative assessment of how geometrical constraint induces the unique microtissue behaviour, and a relevant physics-based model composed of parameters explicit enough to compare with the experimental conditions.

In pursuit of a model-based scaffold design in tissue engineering, we here aim to establish a new framework for simulating complex microtissue deformation processes, including rupture and detachment on structured surfaces, by combining experimental and computational approaches. First, we experimentally investigate the basic features of collective cellular detachment and aggregation *i.e.*, how these actions are influenced by substrate curvature, cellular contractility, and cell-substrate adhesion strength. Second, we propose a new particle-based model that explicitly depicts the process in which cellular contractile forces break cell-cell and cell-substrate adhesions and how this process leads to subsequent ruptures, topological changes, and large deformations of microtissue under three-dimensional geometrical constraints. Finally, based on computational simulations using the developed model and relevant experiments, we clarify the mechanism through which cell-generated forces cause stochastic and catastrophic microtissue changes under 3D constraints. Our physics-based model offers a quantitative and practical methodology for predicting the complex microtissue morphology that cells form on structured surfaces. It could, therefore, serve as a basis for scaffold design in tissue engineering in the future.

## 2. Materials and methods

### 2.1. Cell culture

Human aortic smooth muscle cells (SMCs, KS-4009, Kurabo, Osaka, Japan), a model cell type intrinsically forming a smoothly curved vascular wall *in vivo*, were purchased at passage 3. The cells were maintained in a 100 mm cell culture dish using SmGM2 Smooth Muscle Cell Growth Medium BulletKit (SmGM2, CC-3182, Lonza, Basel, Switzerland) at 37 °C in a humidified atmosphere with 5% CO_2_. The medium was replaced every 3 days. After reaching pre-confluency, the SMCs were detached from the cell culture dish using 1 mL 0.05% Trypsin-EDTA (2530054, Invitrogen, Carlsbad, CA, USA). The cell suspension was then added to 5 mL SmGM2 medium, followed by centrifugation at 1200 rpm for 5 min. The centrifuged cells were suspended again in the medium at defined concentrations for either subculture or each experiment described in the following sections. Cells between passages 6 and 10 were used in this study. To check the homogeneity of SMCs, we cultured them on a glass-bottom cell-culture dish (3911- 035, AGC Techno Glass, Haibara, Japan) for 50 hours and stained α-smooth muscle actin (αSMA), a contractile phenotype marker, using mouse monoclonal anti-αSMA antibody (1:400, A2547, Sigma-Aldrich, St. Louis, MO, USA) and Alexa Fluor 488-conjugated goat anti-mouse antibody (1:200, A11029, Invitrogen), together with nuclear staining using Hoechst 33342 (1:2000, H3570, Invitrogen). Fluorescence microscopy revealed almost homogeneous expression of αSMA in SMCs (Fig. S1). We thus treat SMCs as a homogeneous population in terms of cellular phenotype and contractility in this study.

### 2.2. Microfabricated substrates

To characterise the cellular behaviours on smoothly curved surfaces, custom-made quartz-glass substrates (Shin-etsu Chemical, Tokyo, Japan) were prepared. These substrates have round-ended-trough-shaped, semicircular cross-section concavities at different scales (Fig. 1A); the length-to-width aspect ratio of all the concavities is 5. The scale of the concavities is solely characterised by the diameters of the semicircular cross section (*D* = 200, 300, 400, 500, 600, 1000 µm), which is identical to the concavity widths. We here adopt the curvature of the arc of the concavity cross-section *κ* = 2/*D* as a representative parameter of the geometrical constraints. Note that these trough-shaped concavities have locally different mean curvature *H*, defined as follows:

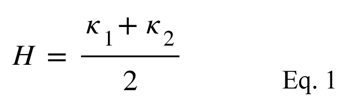

**Fig. 1.**
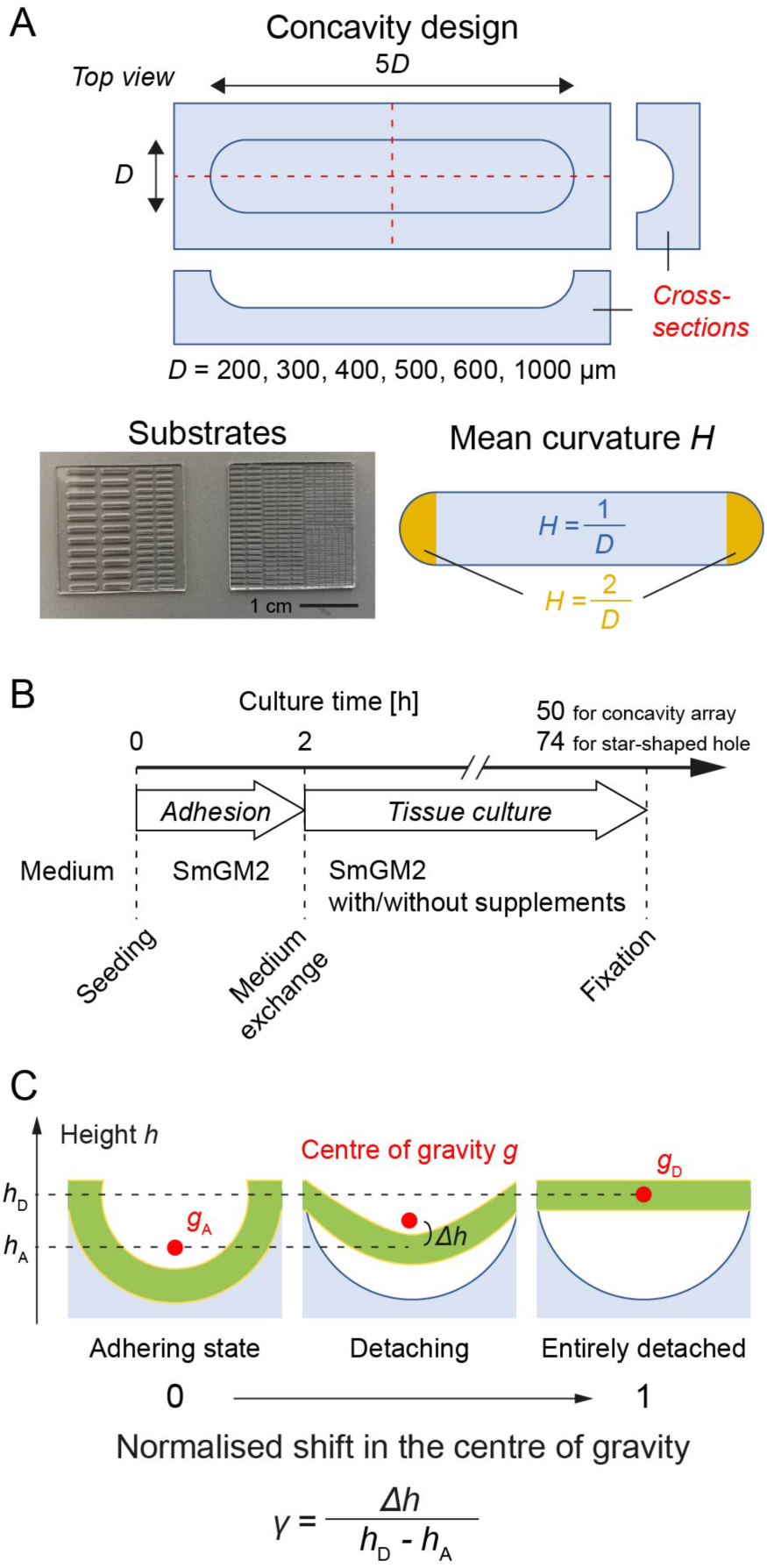
Schematics of the experimental system. (A) Geometry of trough-shaped concavities and substrate pictures. (B) Cell culture protocol. Following a 2 h initial adhesion period, human aortic SMCs were cultured for 48 and 72 h, on the trough-shaped concavities and star-shaped holes, respectively. (C) Schematic of the normalised shift in the centre of gravity *γ* used as an indicator of detachment from curved surfaces. The position of the centre of gravity of the microtissue *g* was calculated from the stack of confocal images. *γ* was subsequently calculated by normalising its height (*i.e.*, *z* position) by those of the adhering cell sheet *h*_A_ and entirely detached one *h*_D_ to quantify the degree of microtissue detachment.

Principal curvatures *κ*_1_ and *κ*_2_ are, respectively, the maximal and minimal curvatures of the surface contour at the given point.

To observe microtissue morphology formed on complex structures, substrates containing star-shaped holes were fabricated via conventional soft lithography. A positive master mould containing microstructures of twelve-cornered stars of different widths (1000 and 2000µm) and corner angles (30°, 60°, 90° and 120°) were fabricated via photolithography using SU-8 2050 (MicroChem, Newton, MA, USA). The height of all the star structures was 100µm. Polydimethylsiloxane (PDMS, Sylgard 184, Dow Chemical, Midland, MI, USA) was poured onto the master mould, cured and removed from it to obtain a silicone elastomer substrate with star-shaped holes.

### 2.3. Surface coating of the substrates

Prior to cell culture, ECM protein was coated onto the microfabricated substrates with or without covalent cross-linking, according to previously published protocols [46] with slight modifications. Either collagen type I (354249, Corning, NY, USA) or fibronectin (F1141, Sigma-Aldrich) was chosen for coating in each experiment. For covalent coating, the microfabricated substrate was rinsed with distilled water, followed by a plasma treatment (RF-Level: High, 1 min, PDC-001, Harrick Plasma, Ithaca, NY, USA). The substrate was sealed in a Teflon box with a microtube containing 3 µL methacryloxypropyltrimethoxysilane (LS-3380, Shin-Etsu Chemical). The box was then kept at 65°C for 2 h. After removing the microtube, the box was baked again for 2 h to complete the silanisation. The substrate was then covered by 1 mM sulfosuccinimidyl 6-(4’-azido-2’-nitrophenylamino)hexanoate (Sulfo-SANPAH, 518-96941, Wako, Osaka, Japan) diluted in PBS (pH = 8.5) and exposed to UV at 40 mW/cm^2^ for 4.5 min, using 100 W Xe lamp (LAX-103, Asahi Spectra, Tokyo, Japan). UV illumination was once repeated after replacing the Sulfo-SANPAH solution. The substrate was immersed in either 0.3 mg/mL Rat tail Collagen Type I diluted in distilled water or 50 µg/mL fibronectin diluted in PBS and kept at 4 °C overnight. The collagen-coated substrate was rinsed with 0.1 M NaOH. The fibronectin-coated substrate was, alternatively, rinsed with distilled water. Following sequential washes with 70% ethanol and PBS, the protein-coated substrates were stored in PBS at 4 °C for cell culture experiments. For non-covalent coating, alternatively, the microfabricated substrate was rinsed with distilled water, treated with plasma (RF-Level: Hi, 1min), and immersed in 0.1 mg/mL poly-D-lysine (PDL, P7886, Sigma-Aldrich) diluted in PBS for 30 min. After washing with distilled water, the substrate was kept in either 0.3 mg/mL collagen I or 50 µg/mL fibronectin solution at 4 °C overnight. The substrate was rinsed, sterilised, then preserved as described above.

### 2.4. Cell-culture procedure on microfabricated substrates

SMCs were cultured on the microfabricated substrates (Fig. 1B). The ECM-coated substrates were preincubated with SmGM2 for 30 min. Suspended SMCs were seeded onto the substrate at 7×10^4^ cells/cm^2^. The cell-seeded substrate was then placed in an incubator for 2 h to allow SMCs to adhere to the substrate surface. Next, we tuned the cellular contractility by replacing the medium with a new one containing recombinant human TGF-β (2.5 ng/mL, 100-21, PeproTech, Rocky Hill, NJ, USA), to promote cellular contractility, or blebbistatin (10µM, 13013, Cayman Chemical, Ann Arbor, MI, USA), to reduce it. Alternatively, we weakened N-cadherin-mediated cell-cell adhesion of SMCs [49] by using the medium with the addition of mouse monoclonal anti-N-cadherin antibody (10 µg/mL, C3865, Sigma-Aldrich). The SMCs were cultured for 48 h on trough-shaped concavity arrays. Meanwhile, SMCs seeded on the substrate with star-shaped holes were cultured for 72 h because they require one additional day to form a confluent sheet covering all the side walls of the holes. Time-lapse imaging was performed using a phase contrast microscope (Eclipse Ti, Nikon, Tokyo, Japan) equipped with a stage incubation system (GM-8000 and WSKMOR-GI, TOKAI HIT, Fujinomiya, Japan), as necessary. The cells were fixed using 4% paraformaldehyde (163-20145, Wako), permeabilised with 0.1% Triton X-100 (T9284, Sigma-Aldrich) diluted in PBS. Following a blocking treatment with Block-Ace (UK-B40, DS Pharma Biomedical, Osaka, Japan), the actin cytoskeleton was stained with Alexa488 phalloidin (1:500, A11029, Invitrogen) overnight.

### 2.5. Assessment of microtissue morphologies cultured on concavity arrays

The fixed SMCs on concavity arrays were observed using a phase-contrast microscope (Eclipse TE300, Nikon). The number of concavities containing SMCs detaching from the curved bottom either partially or entirely, and those forming a clump larger than 100 µm on the substrate surface was counted. Based on this observation, the probabilities of SMC cultures adhering, partially detaching, entirely detaching and aggregating on concavities were calculated by normalising the counts by the total number of the concavities.

### 2.6. Quantitative assessment of cellular detachment from structured substrates

The samples were observed using a confocal laser scanning microscope (LSM700, Carl Zeiss, Oberkochen, Germany) using a 10x lens (N.A. 0.3). The concavities with a diameter of 1000 µm, 600 µm and others were observed with *z*-stepping of 10 µm, 8 µm and 5 µm, respectively. The star-shaped holes were observed with *z*-stepping of 10 µm. From the stack of the confocal images of the actin cytoskeleton, we first extracted the substrate shape to estimate the heights *h_A_* and *h_D_* of the centre of gravity of a microtissue covering the concavity *g_D_* and that of a completely detached microtissue *g_D_*, respectively (Fig. 1C). The centre of gravity *g* of the microtissue inside the concavity was then determined based on the spatial distribution of the binarised actin cytoskeleton to evaluate its vertical shift #ℎ from ℎ_!_. In the end, the normalised shift in the centre of gravity *γ* was quantified as follows:

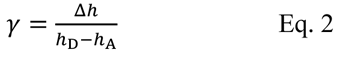

*γ* is 0 when SMCs form a confluent monolayer on the substrate, increases as the detachment progresses, and reaches 1 when SMCs entirely detach from the concavity and form a bridge-like structure connecting both edges of the concavity. To assess the curvature-dependency of microtissue detachment, eight to ten concavities at each radius size were randomly picked, and the *γ* of each concavity was calculated. The median of these *γ* values was treated as the value representing the experiment. In the case of the star-shaped holes, we focused on each triangular corner of the stars as a region of interest, and simply evaluated the shift in height of the centre of gravity of the microtissue #ℎ as an indicator of the degree of detachment. The confocal images were visualised using a custom MATLAB script (2019b, Mathworks, Natick, MA, USA). The data was visualised using either a MATLAB script or Imaris Viewer (Oxford Instruments, Abingdon, UK).

### 2.7. Particle-based microtissue model

To simulate microtissue deformations, we propose a particle-based model that describes three-dimensional dynamics of microtissues at a single cell resolution (Fig. 2A). In this model, individual cells are represented by particles. Adhesions between cells and the substrate are described by the bonds that connect adjacent particles, or that connect particles to the substrate. Microtissue structure is, therefore, expressed by particle locations and the bond network topology, and microtissue deformations are expressed by the motions of cellular particles and the rearrangement of adhesion bonds.

**Fig. 2.**
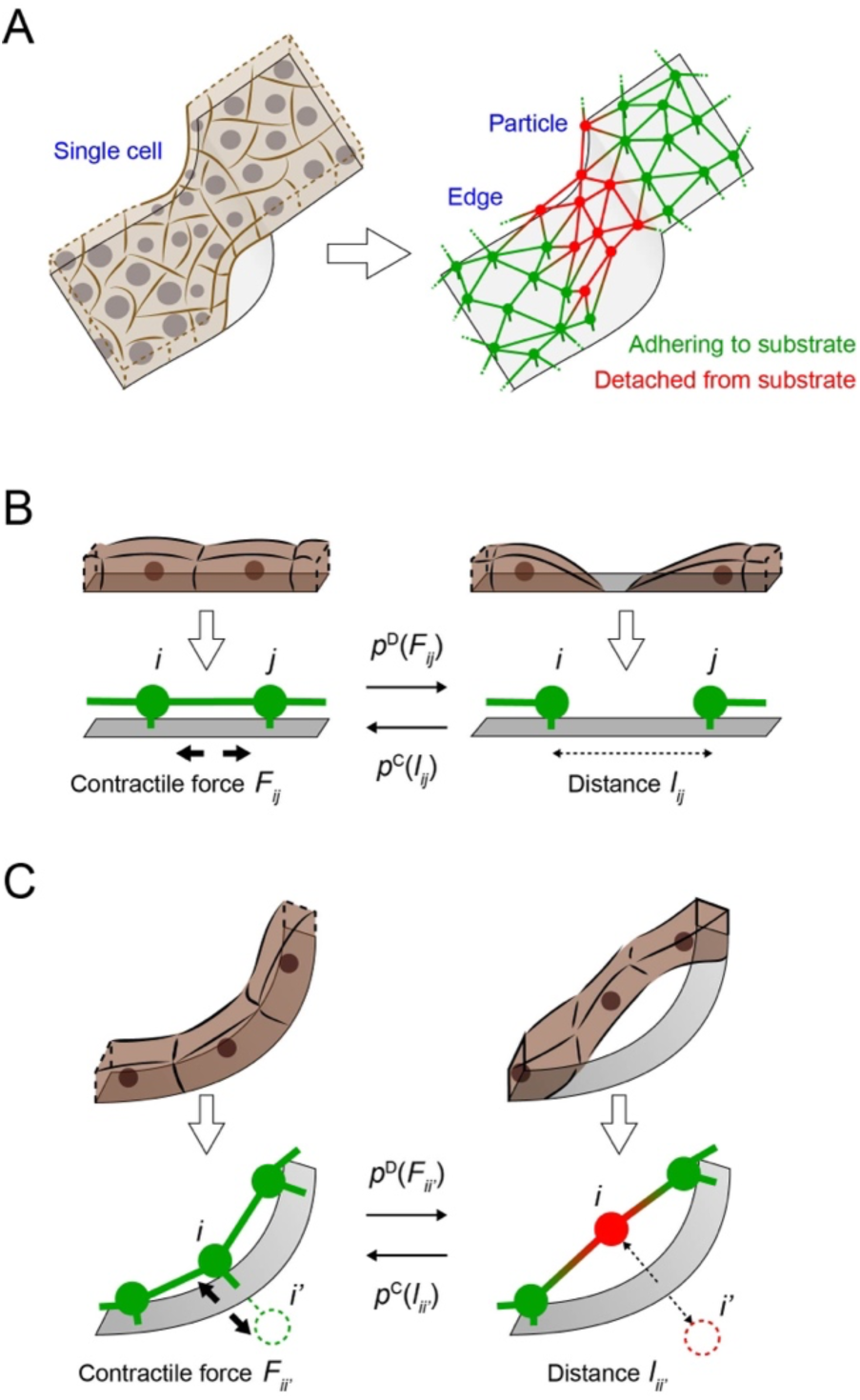
Schematics of the particle-based microtissue model. (A) Microtissue is expressed as a network of particles connected to each other. Cells are expressed as particles, whereas cell-cell and cell-substrate adhesions are expressed as edges. (B) Bonds connecting cellular particles are stochastically formed and eliminated, according to the cell-cell distance and the force applied to the bond connecting the two cellular particles, respectively. (C) The same operation on stochastic formation and elimination is applied to cell-substrate bonds, by considering a mirror boundary condition. The interaction between the *i*-th cell and the substrate is translated to that between the cell *i* and a pseudo cell *i*’ positioned at the mirrored position across the substrate surface.

#### 2.7.1. Description of cell movements and mechanical behaviours

Because microtissues gradually deform at the time scale of hours, microtissue geometry has a stationary and stable instantaneous cell configuration, which satisfies a mechanical force balance [50]. The *i*-th cell location, represented by ***r****i*, is thus obtained by local minimisation of an effective energy function, represented by *U*.

Cells form bonds to each other and to a substrate; these bonds propagate mechanical forces that cells generate, leading to collective deformation of microtissues [39]. To express cell-generated forces, the contractile energy of individual bonds was included. At the same time, the excluded volume effect of individual cells was introduced. Taking these interactions, the effective energy function, *U*, is described as

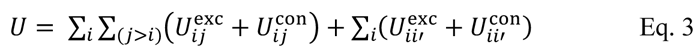

In Eq. 3, the first term indicates cell-cell interactions, whereas the second term indicates cell-substrate interactions. Here, *Uij*^exc^ and *Uij*^con^ are functions of excluded volume energy and contractile energy between the *i*-th and *j*-th cells, respectively. In the second term, cell-substrate interactions were introduced by considering a mirror boundary condition, where the interaction between the *i*-th cell and substrate was handled as that between the *i*-th cell and the corresponding pseudo cell, represented by *i*’, located at the mirrored position across the substrate surface.

The excluded volume energy repels cells from each other. Representing the distance between the

*i*-th and *j*-th cells by *lij*, the excluded volume energy, *Uij*^exc^, is described as

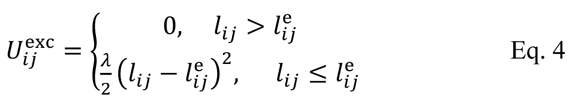

where constants *λ*, and *l*^e^*ij* are a repulsive modulus and a threshold distance, respectively. Constant *l*^e^*ij* is described as the sum of the *i*-th and *j*-th cell radius, *i.e.*, *l*^e^*_ij_* = *σ_i_* + *σ _j_*, where *σ_i_* is the *i*-th cell radius.

The contractile energy attracts cells to each other through their adhesions, whose amplitude is regulated by actomyosin activity inside the cells. The contractile energy, *Uij*^con^, is described as

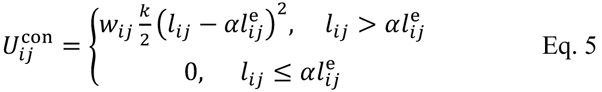

where *k* is a spring modulus. Constant *α*, the amplitude modulus of cell contraction, is a key parameter affecting microtissue morphology. Note that large *α* indicates low contractility and small *α* indicates high contractility, corresponding to treating cells with blebbistatin and TGF-β in the experiments, respectively. Function *wij* is a binary function, which is 1 if the *i*-th and *j*-th cells are connected by an adhesion bond, and 0 otherwise. The same rule was applied to the adhesion bond between the *i*-th cell and the substrate surface (*i.e.*, the corresponding pseudo cell *i*’ considered for the mirror boundary condition).

Cell shapes drastically differ in terms of whether cells attach to the substrate or not, *i.e.*, cells spread out rather flatly when they adhere to the substrate and round out when they leave the substrate.

Therefore, the *i*-th cell radius, *σ_i_*, is described by

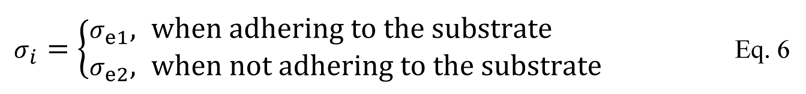

where constants *σ*_e1_ and *σ*_e2_ indicate cell radii when adhering to and separated from the substrate, respectively.

#### 2.7.2. Description of adhesion and separation between cells and substrate

During microtissue deformation, cells dynamically rearrange their adhesions to each other and to the substrate. The formation and rupture of cell-cell and cell-substrate adhesions are expressed in the models by stochastically connecting and disconnecting cellular particles and the substrate. Because adhesive cells form an adhesion to adjacent cells or the substrate when they are in contact, the frequency of adhesion naturally depends on the cell geometry, *i.e.*, distance between cells. On the other hand, the frequency of separation depends on the force acting on the cell-cell boundary, deriving from the mechanosensitive cell-adhesion and cytoskeletal molecules as experimentally shown [51]. Thus, the operations of connecting and disconnecting cells (Fig. 2B) and the substrate (Fig. 2C) are stochastically applied according to their distance and the tension applied to the bond, respectively. The time scale of these adhesion and rupture events is in principle correlated with those of cellular deformation at each interface because these events follow individual cell deformation. Our model, describing each cell as a particle, ignores such microscopic deformation at the interfaces to focus on the macroscopic morphologies of cell clusters observed in the experiments. Instead, the influence of the cell deformation on the cell-cell and cell-substrate adhesion is incorporated into the probability functions defining the frequency of the topological change of the bond network. Moreover, we further simplify the functions by introducing *τ*, a common representative time for both adhesion and separation approximated based on the migration-based remodelling of cell-substrate adhesion. The probability functions of adhesion and rupture of cells and the substrate are thereby expressed as one-variable functions of distance and force, respectively, as described below.

A cell adheres to another cell as well as to the substrate when they physically meet. To determine the probability of forming adhesions, we consider the spatial distribution of individual cell volumes. By employing a free polymer chain model [52], a probability density of the *i*-th cell volume distribution, represented by *ρ_i_* (***r***), is approximately described as the Gaussian distribution as follows, using *σ*_e1_ as a characteristic length of cell volume distributions.

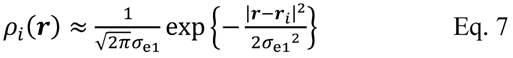

We here consider that two separate cells positioned at ***r****_i_* and ***_r_****_j_* have a chance to contact each other and form a new adhesion. The contact probability at a position ***r*** is considered proportional to the overlap of the spatial distributions of the two cell volumes, whereas the probability with which two cells in contact form a new bond within @ is represented by *ε*(@). Their product is thereby integrated over the entire ***r*** to derive the following bond-formation probability between the *i*-th and *j*-th cells, represented by *a_ij_*, as a function of ***r****_i_*, ***r****_j_* and *t*.

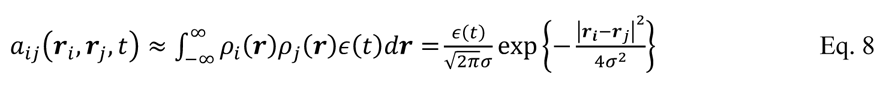

In the end, the probability with which a new bond is formed between the *i*-th and *j*-th cells within a fixed interval G, represented by *p*^C^*ij*, can be derived as follows.

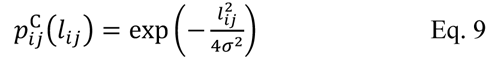

where *l_ij_* = |***r****_i_*-***r****_j_*|. Note that *p*^C^*_ij_* is normalised to satisfy *p*^C^*_ij_* (0) = 1 and *p*^C^*_ij_* (∞) = 0, *i.e.*, the *i*-th and *j*-th cells, always form the adhesion bond within in the limit of *l_ij_* → 0, whereas they do not form a new bond in the limit of *l_ij_* → ∞.

We next define the probability with which cell-cell and cell-substrate adhesions rupture. An experimental study reported a sigmoidal relationship between the survival rate of cell adhesion and the shear stress applied in a certain period [51]. Based on this study, the probability, *pij*^D^, with which the bond between the *i*-and *j*-th cells is dissected within G can be defined as a function of the bond contraction force represented by *F_ij_* (= -∂*U_ij_*/ ∂*l_ij_*), as follows.

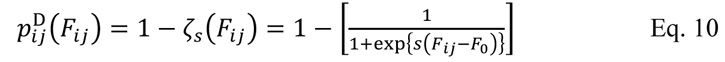

where *ζ_s_* is a normalised logistic function with slope *s*. In the limit of *F_&’_* → +∞, the bond between the *i*-th and *j*-th cells always collapses. Constant *F*_0_ represents the midpoint value of the logistic function, indicating the rupture strength of bonds. Because cells form different molecular complexes at cell-cell and cell-substrate adhesion interfaces [53], these adhesions naturally have different separation patterns at a single cell scale [54]. Observation of a cell cluster on a flat substrate also indicated that the balance of cell-cell and cell-substrate affinity determines whether cells form a monolayer sheet or an aggregated clump [55], highlighting the importance of clearly distinguishing cell-cell and cell-substrate adhesion rupture strengths. Hence, we introduce two distinct rupture strengths for cell-cell and cell-substrate bonds as

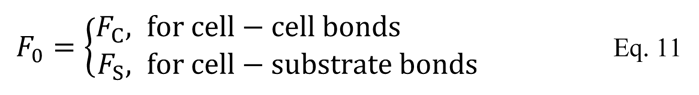

where *F*_C_ and *F*_S_ represent the strengths of cell-cell and cell-substrate bonds, respectively.

*F*_S_ was swept as a key parameter in the simulation, in correspondence with the experimental tuning of cell-substrate adhesion strength using physisorbed collagen (PDL-collagen, weaker adhesion) and chemically crosslinked collagen (SS-collagen, stronger adhesion). Contribution of *F*C to cellular detachment from the substrate was also verified, in correspondence with the experimental tuning of cell-cell adhesion using anti N-cadherin antibody (weaker adhesion).

#### 2.7.3. Nondimensionalisation and parameter values

The physical parameters in the proposed model were nondimensionalised by introducing unit values, which are given by experimental measurements. Unit length, corresponding to cellular size 27_(4_, is set to 41 µm, which is the estimated mean distance between the SMCs confluently cultured at the number density of 7×10^4^ cells/cm^2^. Unit energy is set to the elastic modulus of a SMC, which is *k* = 30 nN/µm. This value was estimated from the slope of the force-strain curve of a single SMC reported in literature [56]. The measured contractile force of a single SMC of 0.2 µN was divided by a strain of 10% and an approx. 70 µm axial length of the stretched SMC to estimate the magnitude of the representative cellular contractility. The representative time of rearranging cell-cell and cell-substrate bonds is roughly estimated from that of the typical size and motility of SMCs; *i.e.*, based a reported migration speed of 12−16 µm/h for vascular SMCs [57] and the unit cellular length 2*σ*_e1_ = 41 µm, the time scale of bond network rearrangements can be estimated to be on the order of hours. We thus defined the unit time G = 4.8 h to track the relaxation-based morphological change in the microtissue for 48 h within 10 steps of bond network rearrangements. Hereafter, individual parameter values are represented using dimensionless values.

The volume of a single SMC of 2×10^4^ µm^3^ was estimated from the measured number density of 7×10^4^ cells/cm^2^ and the height of a confluent SMC monolayer of 6 µm measured via confocal microscopy. Keeping the cellular volume unchanged throughout the detachment from the solid substrate surface, the diameter of the floating cells is set to *σ*_e2_ = 3 µm (= 0.6*σ*_e1_).

To calculate a stationary and stable cell configuration, local minima of the effective energy function were given by solving

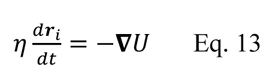

where *n* is the effective friction coefficient (VX = V/ZG = 1). The time integration of Eq. 13 was solved using the Euler method with a time step of *ṫ* = 0.04. The cell locations reached the equilibrium state within G, the time interval for rearranging the bond network between cells and substrate; therefore, the time scale of microtissue dynamics is governed by G°. Note that V is only used for minimising the total energy and does not represent a real physical quantity.

In this present work, we aim to reveal how the interplay of cellular contractility and cell-substrate adhesion strength contributes to the dynamics of microtissue deformation under geometrical constraints in three dimensions. We thus chose the amplitude modulus of cell contraction *α*, the rupture strength of cell-substrate bonds *F*_S_, and that of cell-cell bonds *F*_C_ as key parameters, and varied them to express the effect of drug treatment, surface coating and antibody treatment, respectively. In the standard condition, we set *F*_S_ = 0.5 and *F*_C_ = 0.75, *i.e.*, cell-cell adhesion has 1.5 times larger rupture strength than cell-substrate adhesion, to reproduce the tendency that SMCs detach from structured surfaces. The parameters included in the model are summarised in Table 1. For the detailed values assigned to *α*, *F*_C_ and *F*_S_ in each simulation, we direct readers to Table S1 in the supplementary information.

**Table 1.** List of parameters

| Parameter | Description | Value | Normalisation | Note |
| --- | --- | --- | --- | --- |
| $\sigma_{e1}$ | Size-exclusion radius of a cell adhering to the substrate | 21 $\mu\text{m}$ | $\tilde{\sigma}_{e1} = 0.5$ | Half value of the mean cell-cell distance estimated from measured cell density $7 \times 10^4$ cells/$\text{cm}^2$ at confluency |
| $\sigma_{e2}$ | Size-exclusion radius of a cell detached from the substrate | 13 $\mu\text{m}$ | $\tilde{\sigma}_{e2} = \sigma_{e2}/2\sigma_{e1} = 0.3$ | Estimated from measured cell density $7 \times 10^4$ cells/$\text{cm}^2$ and cell height 6 $\mu\text{m}$ at confluency |
| $\lambda$ | Repulsive modulus of cellular excluded volume | 300 nN/$\mu\text{m}$ | $\tilde{\lambda} = \lambda/k = 10$ | Premodulated |
| $k$ | Elastic modulus of cells | 30 nN/$\mu\text{m}$ | $\tilde{k} = 1$ | Estimated from literature [56] |
| $\alpha$ | Amplitude modulus of cell contraction | - | - | Varied |
| $F_C$ | Rupture strength of cell-cell bonds | - | $\tilde{F}_C = F_C/2k\sigma_{e1}$ | Varied |
| $F_S$ | Rupture strength of cell-substrate bonds | - | $\tilde{F}_S = F_S/2k\sigma_{e1}$ | Varied |
| $\tau$ | Time interval to rearrange the connections | 4.8 h | $\tilde{\tau} = 1$ | Defined based on the reported migration speed of vascular SMCs [57] and $\sigma_{e1}$ |
| $\Delta t$ | Time step for iterations in Euler method | - | $\Delta\tilde{t} = \Delta t/\tau = 0.04$ | Sufficiently smaller than $\tau$ to find the local energy minima |
| $\eta$ | Effective friction coefficient | - | $\tilde{\eta} = \eta/k\tau = 1$ | Optimised for numerical simulations |
| $s$ | Steepness of the logistic function included in $p_{ij}^D$ | 12.5 | - | Optimised for allowing cells to form a stable monolayer sheet at the initial state of simulation |

#### 2.7.4. Boundary condition, initial condition, and simulation procedures

A flat square plane containing a concavity of interest in the centre was designed to represent a substrate surface. Trough-shaped concavities and star-shaped holes of different scales, identical to the fabricated substrate shapes, were used in this study. The inner wall of a smoothly bending tube, which has an identical shape to a hollow hydrogel tube reported in a previous study [47], was also adopted as a boundary condition to predict the microtissue deformation in confined spaces. Periodic boundary conditions were imposed in both the *x-* and *y-* directions, and the solid boundary condition with zero restitution coefficient was imposed on the substrate. In experiments, cells typically form a monolayer sheet on structured surfaces at the initial stage of the cell culture, regardless of whether they stably keep the structure afterwards or not. The initial condition was therefore created via the following process: First, cells were randomly located in the plane on the substrate at the density equivalent to a confluent cell sheet. Second, cell positions were equilibrized by the cell motions according to the excluded volume energy of Eq. 4 while keeping the distance of each cell to the substrate at a constant value sufficiently small to form a cell-substrate bond. Using this initial condition, numerical simulations were performed within 10G° (*i.e.*, 48 h), unless otherwise noted, as considering both the particle motions governed by the total energy of Eq. 3 and the bond network rearrangements. The framework was implemented in C++. The simulation was run on a 4-core Intel Xeon E-2224 3.4 GHz or a 6-core Intel Core i7-8700B machine equipped with 64 GB RAM. The numerical simulation within 20G° using the trough-shaped concavity surface as a boundary condition takes approximately 4 min for *D* = 200 µm (505 particles), 26 min for *D* = 500 µm (3160 particles), and 280 min for *D* = 1000 µm (12641 particles). The results were visualised using ParaView ver. 5. 9. 1b [58].

### 2.8. Statistical analysis

The cell culture experiments and the simulations were at least triplicated to confirm the reproducibility of the results. The error bars shown in the figures represent either the standard errors (SE) or the standard deviations (SD) of the experiments and simulations repeated independently, as indicated in each figure caption with repeat number *N*.

## 3. Results

### 3.1. Substrate curvature, cellular contractility and ECM-substrate adhesion strength stochastically influence microtissue morphology on concavity

Conventional studies have experimentally shown that cellular detachment [45–47] and aggregation [46,48] are induced by surface curvature and cell-generated force, the mechanism has yet to be elucidated. To clarify the mechanical features of cellular detachment, we first cultured SMCs on trough-shaped concavity arrays, varying the surface curvature of the substrate, the adhesion strength of ECM to the substrate, the cellular contractility, and the strength of N-cadherin-mediated cell-cell adhesion.

Time-lapse imaging of SMCs cultured on a concavity of 1000 µm diameter in the presence of TGF-β revealed a typical microtissue deformation process on curved surfaces (Fig. 3A and supplementary movie 1). SMCs first formed a continuous monolayer on the concavity. The cell sheet partially ruptured and detached from the substrate surface at around 48 h of cell culture. The partial rupture quickly propagated to the entire cell sheet due to the shrinkage of the detached cells. The microtissue consequently aggregated into a clump following the successive detachment and shrinkage. During the deformation process, SMCs showed three characteristic forms: a sheet form covering the substrate surface (adhering state), a bridge-like structure formed inside the concavity due to either partial or entire detachment of the cell sheet (partial or entire detachment) and an aggregated clump (aggregation). This observation led us to hypothesise that intrinsic cellular contractile force impaired cell-substrate adhesion, and it initiated the successive compaction of cells, rupture of cell-cell adhesions, and final aggregation on curved surfaces.

**Fig. 3.**
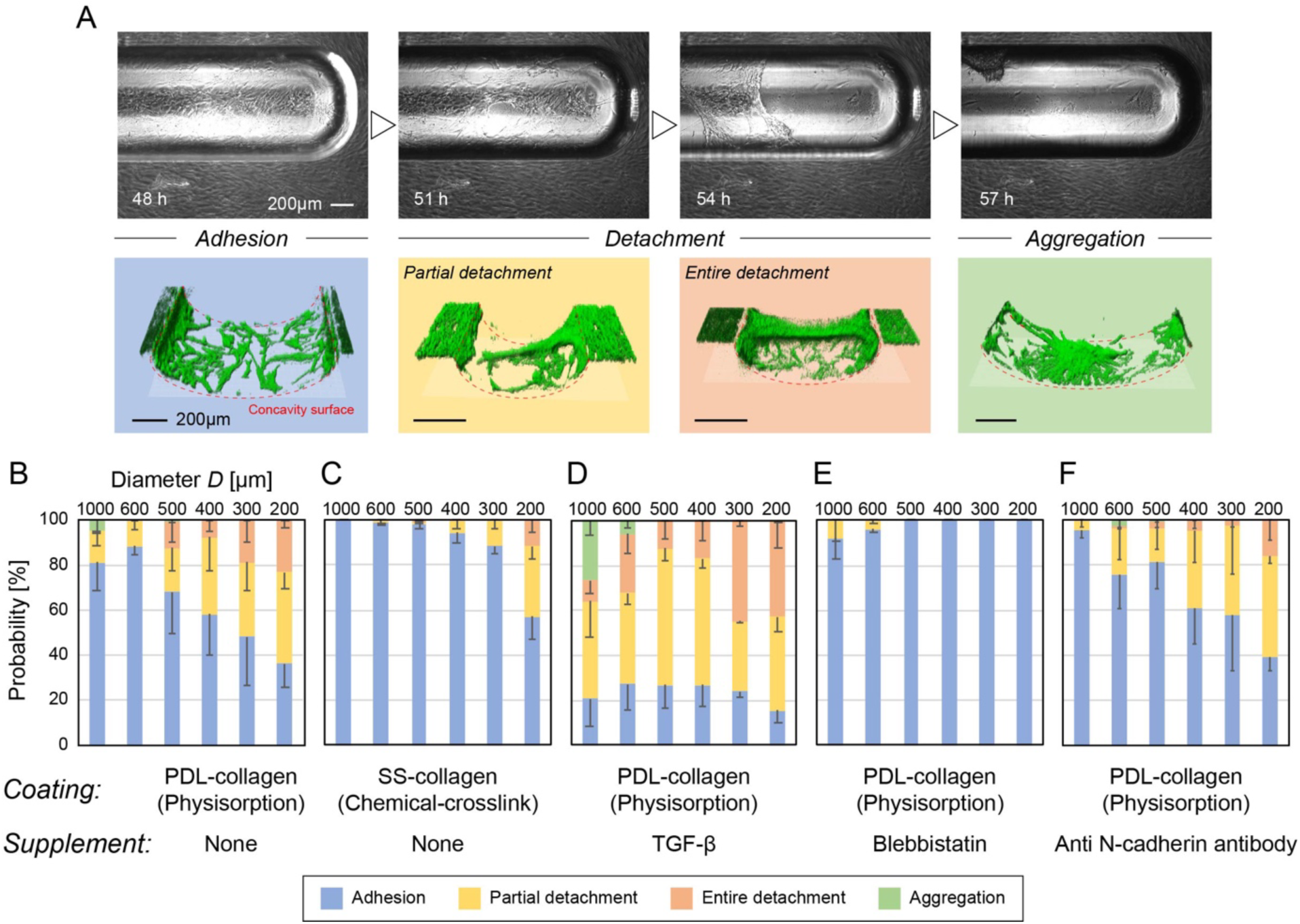
Deformation process and characterisation of morphologies of SMCs cultured on trough-shaped concavity arrays. (A) Representative morphologies of SMCs observed on trough-shaped concavities. The series of time-lapse images show how SMCs spontaneously detach from a collagen-physisorbed concavity surface and aggregate under the presence of TGF-β. Corresponding typical four morphologies, *i.e.*, adhering, detaching either partially or entirely, and aggregating states of SMCs observed via confocal microscopy are listed below. (B–E) Appearance probability maps of the four morphologies of SMCs cultured on the concavity arrays of different diameters (mean-SE, *N* = 3). Collagen was coated onto the substrate via either physisorption using PDL (B, D, E) or chemical crosslink using SulfoSANPAH (C) to tune the adhesion strength of the ECM to the substrate surface. TGF-β (D) and blebbistatin (E), respectively, were added to the medium to promote and suppress cellular contractility. Anti-N-cadherin antibody was added to the medium to suppress the cell-cell adhesion (F).

Next, we systematically investigated the influence of the surface curvature, the adhesion strength of ECM to the substrate, and the cellular contractility, on the microtissue morphology. As a quantitative criterion, we assessed the probabilities with which cells stay in an adhering state, detach either partially or entirely, or aggregate on trough-shaped concavities after 48 h of cell culture. Typical microtissue morphologies observed in each condition are overviewed in Fig. S2. We first assessed the influence of cell-substrate adhesion strength by comparing the morphologies of SMCs on concavities coated with physisorbed and chemically crosslinked collagen. SMCs spontaneously detached from collagen-physisorbed concavities, and the probability of detachment increased as the surface curvature increased (*i.e.*, concavity diameter *D* decreases) (Fig. 3B). Notably, SMCs aggregated only on the concavity of the lowest curvature (*D* = 1000 µm) with several percent of probability. In contrast, SMCs showed considerably smaller detachment probability on the collagen-crosslinked concavities; the curvature dependency nevertheless still obtained (Fig. 3C). Generally, SMCs maintained a stable sheet form on the collagen-crosslinked concavities and did not form aggregated clumps. SMCs on fibronectin-crosslinked concavities also yielded a nearly identical probability map (Fig. S3). These observations clearly indicate that the adhesion strength of ECM onto the scaffold surface is a significant mechanical parameter affecting the microtissue morphology on curved surfaces.

We next added either TGF-β or blebbistatin to the medium to assess the influence of cell-generated force on the morphology of SMCs on concavities. Addition of TGF-β, which promotes contraction of SMCs, considerably elevated the probabilities of either partial or entire detachment from the concavities (Fig. 3D). The probability of the adhering state under these conditions decreased to less than 30%. TGF-β also promoted aggregation of SMCs at low curvatures (*D* = 600, 1000 µm). It was remarkable that net probability of detachment (partial or entire) increased with the decrease in concavity diameter, showing a clear curvature dependency. In contrast, addition of blebbistatin, which attenuates cellular contraction, generally kept SMCs in an adhering state on all concavities (Fig. 3E), though there was still less than 10% probability of partial detachment at smaller curvatures. Antibody treatment, which inhibits N-cadherin-mediated cell-cell adhesion, tended to decrease the probability of entire detachment (Fig. 3F), but it did not have a major influence on the net of probability of detachment (partial or entire). Notably, there were still a few percent of aggregation; this was observed at higher curvatures around 600 µm. These observations clearly demonstrate that collective cellular contraction is a driving factor of morphological change on concavities.

These probability maps indicate that curvature-dependent cellular detachment and aggregation is initiated by a rupture of weak cell-substrate adhesion. The rupture event appears in large part to be a mechanical event driven by cell-generated forces. These observations highlight the importance of substrate curvature, cellular contractility, and cell-substrate adhesion strength as key factors affecting microtissue morphology formed on structured surfaces. It is also notable that drastic deformation of cell sheets on curved surfaces seems to occur in a rather probabilistic way, unlike the deterministic tissue morphogenic process often found *in vivo*. These features suggest that spontaneous microtissue deformation, triggered by a rupture of cell-substrate adhesion, is basically a stochastic event.

### 3.2. Cell-generated force ruptures cell-substrate adhesion and drives curvature-dependent cellular detachment

We next focused on the details of the cellular detachment. SMCs were cultured on the trough-shaped concavities, and the normalised shift in the centre of gravity *γ* (Fig. 1C), defined by Eq. 1, was evaluated as a quantitative readout of cellular detachment. The influence of cell-substrate adhesion strength was first evaluated by comparing *γ* of SMCs cultured on the concavities coated with physisorbed collagen (PDL-collagen), chemically crosslinked collagen (SS-collagen), and chemically crosslinked fibronectin (SS-fibronectin) (Fig. 4A). *γ* generally increased as the surface curvature increased, supporting the idea that the surface curvature causes the spontaneous detachment of cells [45–47]. Chemical crosslink of ECM attenuated *γ*; the influence was noticeable at curvatures larger than 5 mm^-1^. Notably, only a small difference was found between crosslinked fibronectin and collagen, in accordance with the probability maps obtained in the previous morphological observations.

**Fig. 4.**
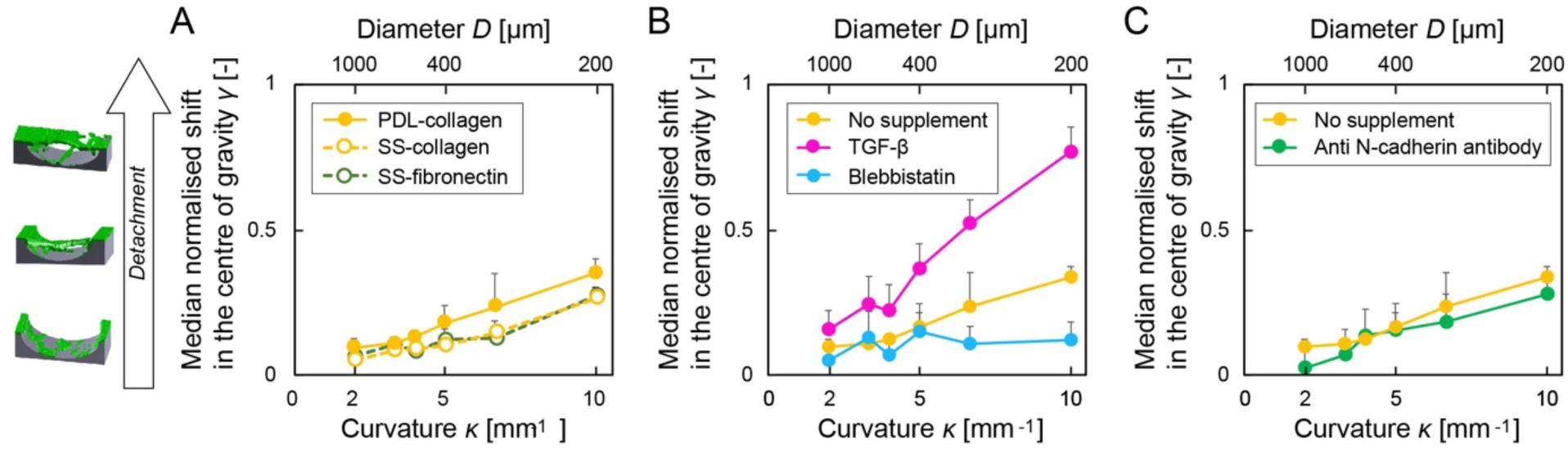
Characteisation of microtissue detachment from trough-shaped concavities. (A) Comparison of the influence of ECM coatings on the microtissue detachment (mean+SD, *N* = 3). Collagen was coated onto the substrate via physisorption using PDL (PDL-collagen). Collagen and fibronectin were coated onto the substrate via chemical crosslink using Sulfo-SANPAH (SS-collagen and SS-fibronectin). (B) Comparison of the influence of cellular contractility on the microtissue detachment (mean+SD, *N* = 3). Collagen was coated onto the substrate via physisorption using PDL. TGF-β and blebbistatin were added to the medium during the cell culture term to promote and suppress cellular contractility, respectively. (C) Comparison of the influence of cell-cell adhesion to the microtissue detachment (mean+SD, *N* = 3). Collagen was coated onto the substrate via physisorption using PDL (PDL-collagen). Anti-N-cadherin antibody was added to the medium to suppress N-cadherin-mediated cell-cell adhesion.

The influence of cellular contractility was next assessed by adding TGF-β and blebbistatin into the medium. The TGF-β treatment promoting cellular contractility elevated *γ*, increasing the curvature dependency. In contrast, the blebbistatin treatment reducing the cellular contractility kept *γ* below 0.1. These contrasting observations support the idea that cell-generated force is a major factor inducing a rupture of cell-substrate adhesion on curved surfaces, in good agreement with the general tendency of the appearance probability maps of microtissue morphologies (Fig. 3B, D and E). The curvature dependency of the spontaneous cellular detachment is also in good agreement with the conventional reports on curvature-induced cellular detachment [45–47], confirming that our concavity array is a useful platform to extensively examine the effect of surface curvature on the cell morphologies. *γ*, interestingly, tended to show more variance at a medium curvature of around 5 mm^-1^ at around 0.2. This tendency was even more evident in the presence of TGF-β. These features imply that the cellular detachment is accelerated by cellular contractility at the initial step, indicating the catastrophic character of the event.

The influence of cell-cell adhesion was also assessed by adding anti-N-cadherin antibody into the medium. The antibody treatment inhibiting N-cadherin-mediated cell-cell adhesion tended to decrease *γ* slightly (Fig. 4C), but SMCs clearly showed detachment from the concavity bottoms at curvatures higher than 5 mm^-1^ (Fig. S2). Overall, the effect of antibody treatment in this experiment was not as large as that of blebbistatin treatment, *i.e.*, inhibition of cellular contractility.

### 3.3. Particle-based model reproduces characteristic deformation process and morphologies of microtissue on concavity

To understand the mechanical basis of drastic microtissue deformation on structured surfaces, we here developed a physics-based model. This model describes a cell as a particle and translates cell-cell and cell-substrate adhesions into bonds between particles to expresses microtissue deformation as a rearrangement of the network of connected particles. The bond network is stochastically rearranged, expressing the stochastic formation and rupture of cell-cell and cell-substrate adhesions during microtissue deformation. Based on the experimental observations, rupture strength of cell-substrate bonds *F*_S_ and amplitude modulus of cell contraction *α* were nominated as key parameters. Applying the trough-shaped concavities as a boundary condition, we simulated the morphological change of a SMC sheet on curved surfaces at single-cell resolution.

The simulation reproduced the process by which cells detach from the substrate under the influence of their own mutually-applied contractile forces (Fig. 5A, supplementary movie 2). Cells initially formed a monolayer sheet of a uniform density on the substrate (*ṫ* = 1). Their contractile interaction induced a condensation of cells mainly on both the edges of the concavities, leading to local compaction and partial detachment from the curved surface (*ṫ* = 5). Detachments then propagated to the central part of the concavity (*ṫ* = 10). Following the successive compaction, cells formed a bridge structure connecting both the edges of the concavity, or a clump adhering close to the edge (*ṫ* = 20). Throughout the simulation, four representative forms of SMCs observed in the experiments were reproduced (Fig. 5B). The correspondence of the predicted microtissue morphologies *in silico* with those observed in the experiment show a certain relevance to explaining the unique microtissue deformations based on the mechanics, *i.e.*, probabilistic adhesion and separation of cells and minimisation of effective energy. We thus utilised the model to investigate the mechanical basis of microtissue deformation on the structured surfaces, as described in the following sections.

**Fig. 5.**
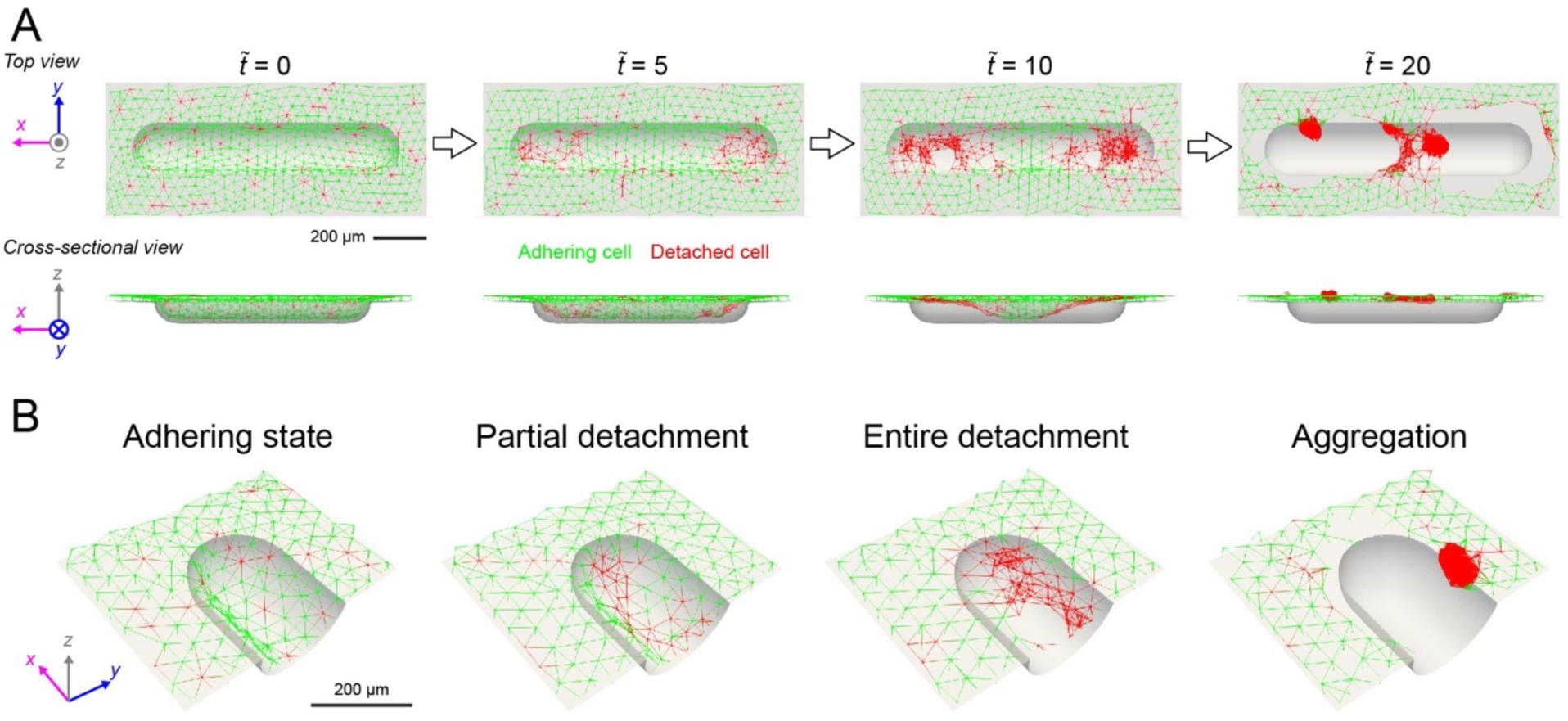
Representative cellular morphologies captured in the simulation. (A) Time course of the simulation. (B) Magnified pictures of the concavity corner, representing the four characteristic morphologies of cellular particles.

### 3.4. Particle-based model predicts influence of cell-substrate and cell-cell adhesion strengths and cellular contractility to catastrophic cell-sheet detachment on concavities

Utilising the particle-based model, we next systematically investigated the quantitative contribution of the substrate curvature *κ*, the rupture strength of cell-substrate adhesion *F*_S_, the amplitude modulus of cell contraction *α*, and the rupture strength of cell-cell adhesion *F*_C_ to the cell-sheet detachment. Sweeping *F*_S_, *α* and *F*_C_ independently, multicellular dynamics on trough-shaped concavities (*D* = 200−1000 µm, covering 2−10 mm^-1^ of substrate curvature *κ*) were simulated. Note that *α* is an indirect indicator of cell contraction, *i.e.*, varying *α* from 0.9 to 0.7 indicates three times larger cell contractility, whereas *F*_S_ directly represents the magnitude of the rupture strength of cell-substrate adhesion. During the simulation, *γ*, the normalised shift in the centre of gravity, monotonically increased over time, indicating the progress of the detachment of cells from the substrate surface (Fig. 6A). The steepness of the chart, reflecting the speed of the detachment, was larger at tighter substrate curvature (*i.e.*, smaller concavity diameter *D*), clearly demonstrating that substrate curvature accelerates the cellular detachment. In contrast to the monotonic relationship between *γ* and *κ*, the standard deviation of *γ* was largest at *D* = 300 µm. This large deviation was attributed to a mixed population of simulation results that showed and did not show cellular detachment within 109°, clearly reflecting the stochastic aspect of the detachment event. At the intermediate curvature, whether cell-substrate bonds ruptured or not was highly probabilistic due to a subtle balance between the net force of cellular contraction and the rupture strength of cell-substrate adhesion.

**Fig. 6.**
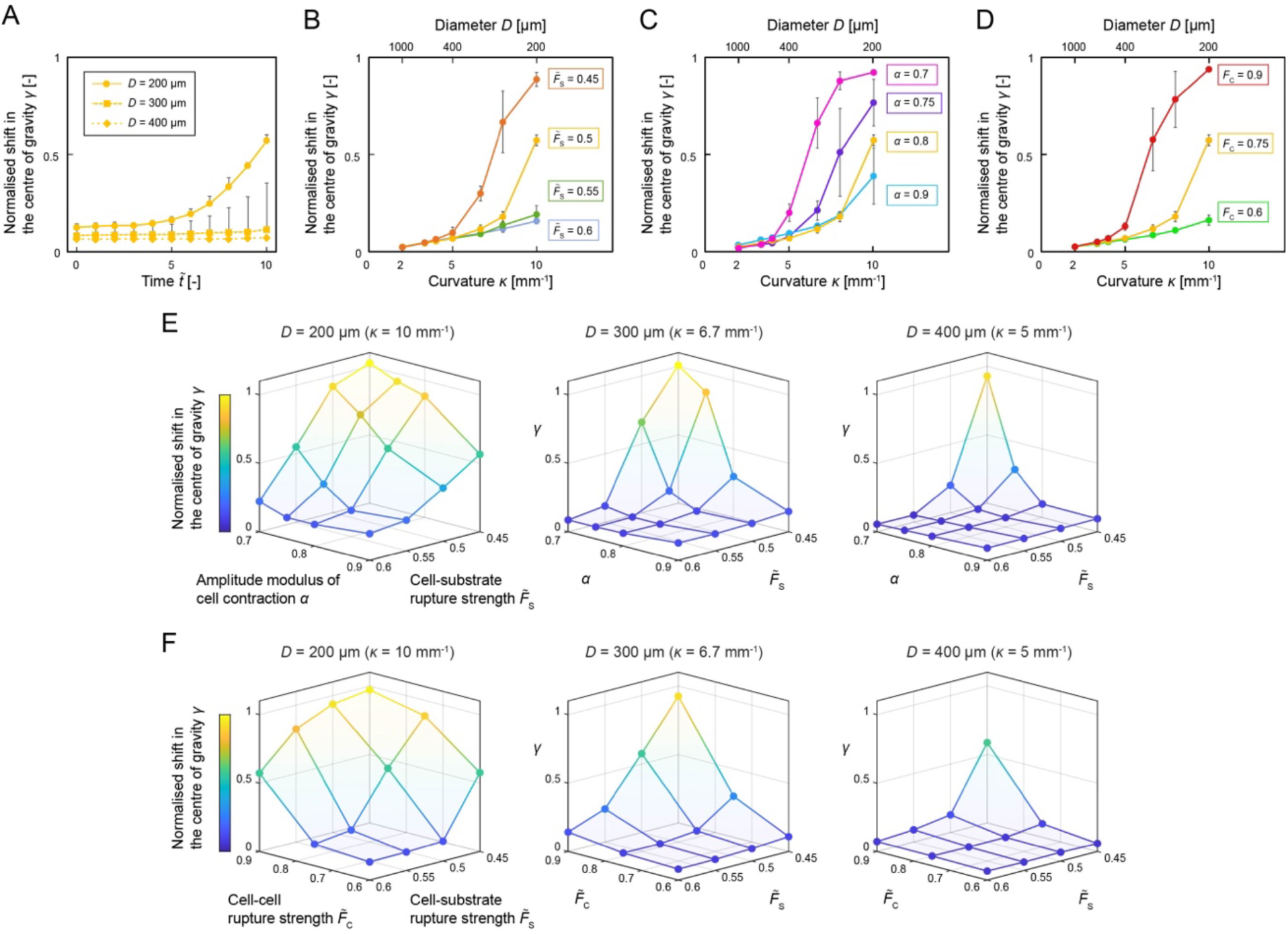
Microtissue detachment simulated by the computational model. (A) Time course of simulated microtissue detachment on concavities with 200, 300 and 400 µm of diameter (mean+SD, *N* = 5). (B) Influence of cell-substrate rupture strength *F*_S_ to the curvature-dependent microtissue detachment at *ṫ* = 10 (mean+SD, *N* = 5). (C) Influence of cell contraction modulus *α* on the curvature-dependent microtissue detachment at *ṫ* = 10 (mean+SD, *N* = 5). (D) Influence of cell-cell rupture strength *F*_C_ to the curvature-dependent microtissue detachment at *ṫ* = 10 (mean+SD, *N* = 5). (E) Comparison of the influence of *F*_S_ and *α* on the microtissue detachment on concavities with different diameters at *ṫ* = 10. (F) Comparison of the influence of *F*_S_ and *F*_C_ on the microtissue detachment on concavities with different diameters at *ṫ* = 10. (E, F) The charts show the means of repeated simulations (*N* = 5).

We then swept each key parameter to predict the influence of cell-substrate as well as cell-cell adhesion strengths and cellular contractility on cellular detachment, by comparing *γ* at *ṫ* = 10, corresponding to 48 h of cell culture time. Decrease in *F*_S_ resulted in a considerable increase in *γ*, changing the curvature dependency from a linear to a sigmoidal one (Fig. 6B). Decrease in *α*, indicating enhanced cellular contractility, also elevated *γ* and made the curvature dependency sigmoidal (Fig. 6C). Increase in *F*_C_ also had a similar influence on *γ*, increasing the sensitivity of detachment to the substrate curvature (Fig. 6D). In all cases, the standard deviation tended to be particularly large at middle curvature around 6 mm^-1^ and at the middle stage of the detachment. These features indicate that the transitional state of the cell sheet during detachment is unstable, providing evidence that the cellular detachment is a non-linear, catastrophic event. The magnitude relationship of *γ* to *F*_S,_ *α* and *F*_C_ was consistent with the experimental results tuning the ECM tethering (Fig. 4A), the cellular contractility (Fig. 4B) and N-cadherin-mediated cell-cell interaction (Fig. 4C), supporting the idea that the particle-based model captures the basic features of the cell-sheet detachment. It should be mentioned, however, that *γ* exhibited a rather linear relationship to the substrate curvature *κ* in the experiments, while the predicted sigmoidal relationship better resembled the experimental outcomes reported previously [46].

The contribution of *F*_S_ and *α* to cell-sheet detachment was next compared at different curvatures (*D* = 200, 300 and 400 µm). (Fig. 6D). Decrease in either *F*_S_ or *α* generally increased *γ*, confirming that cellular contraction impairs cell-substrate adhesion. At the highest curvature (*D* = 200 µm), *γ* showed more sensitivity to change in *F*_S_ than to that of *α* if we compare the slope of the trajectories on *γ* - *F*_S_ and *γ* - *α* planes, but the difference in the sensitivity was less visible as the substrate curvature decreased (*D* = 300 and 400 µm). These results predict that cell-substrate adhesion strength, rather than cellular contractility, has the greater influence on the stability of a cell sheet on structured surfaces, and the relative influence becomes larger as the substrate structure becomes smaller. In addition, our model predicted that cell-sheet detachment occurs when the rupture strength of cell-cell adhesion *F*_C_ is much larger than *F*_S_ (Fig. 6E). This magnitude relationship between *F*_C_ and *F*_S_ indicates that propagation of cellular forces via stable cell-cell adhesion is required to break the cell-substrate adhesion. These results thus highlight the importance of careful tuning of ECM tethering onto the scaffolds as well as the propagation of contractile forces via intercellular connections, when guiding cells to form a monolayer on 3D microstructures.

### 3.5. Stress concentration drives collective cellular detachment from concavity

The mechanical basis of cell-sheet detachment has been unexplored due to the difficulties in measuring invisible cellular forces at complex cell-substrate interfaces. To reveal the detailed mechanical process in which cellular force breaks the cell-substrate adhesion, we utilised our particle-based model and analysed the transition of *F*_bond_, the contractile force applied to each bond, and *F*_wall_, the contractile force that each cell applies to the substrate wall (Fig. 7A, supplementary movie 3). At the early stage of the simulation (*ṫ* = 1), *F*_wall_ applied to the concavity surface (*D* = 200 µm) showed a local increase at both the edges of the trough-shaped concavity, where the mean curvature *Η* is two times higher than the central part of the concavity. At the same time, cells partially detached from the round bottom of the concavity. The tensed cells locally increased *F*_wall_ at the periphery of the partially detaching cell sheet, resulting in a further expansion of the detached area (*ṫ* = 6). The wave of cellular detachment propagated to the central part of the concavity, involving other cells that already had detached (*ṫ* = 10). Local increase in contractile force was also observed in *F*_bond_, especially in the regions close to the concavity edge; this tendency became more and more evident as the detachment progressed. Throughout the deformation process, the simulation clearly demonstrated that a feedback loop of the local increase in *F*_wall_ and *F*_bond_, the break in cell-substrate adhesion, and the shape optimisation of the detached cells accelerated the cell-sheet detachment. As the concavity diameter *D* increased to 400 µm, less detachment was observed (Fig. S4).

**Fig. 7.**
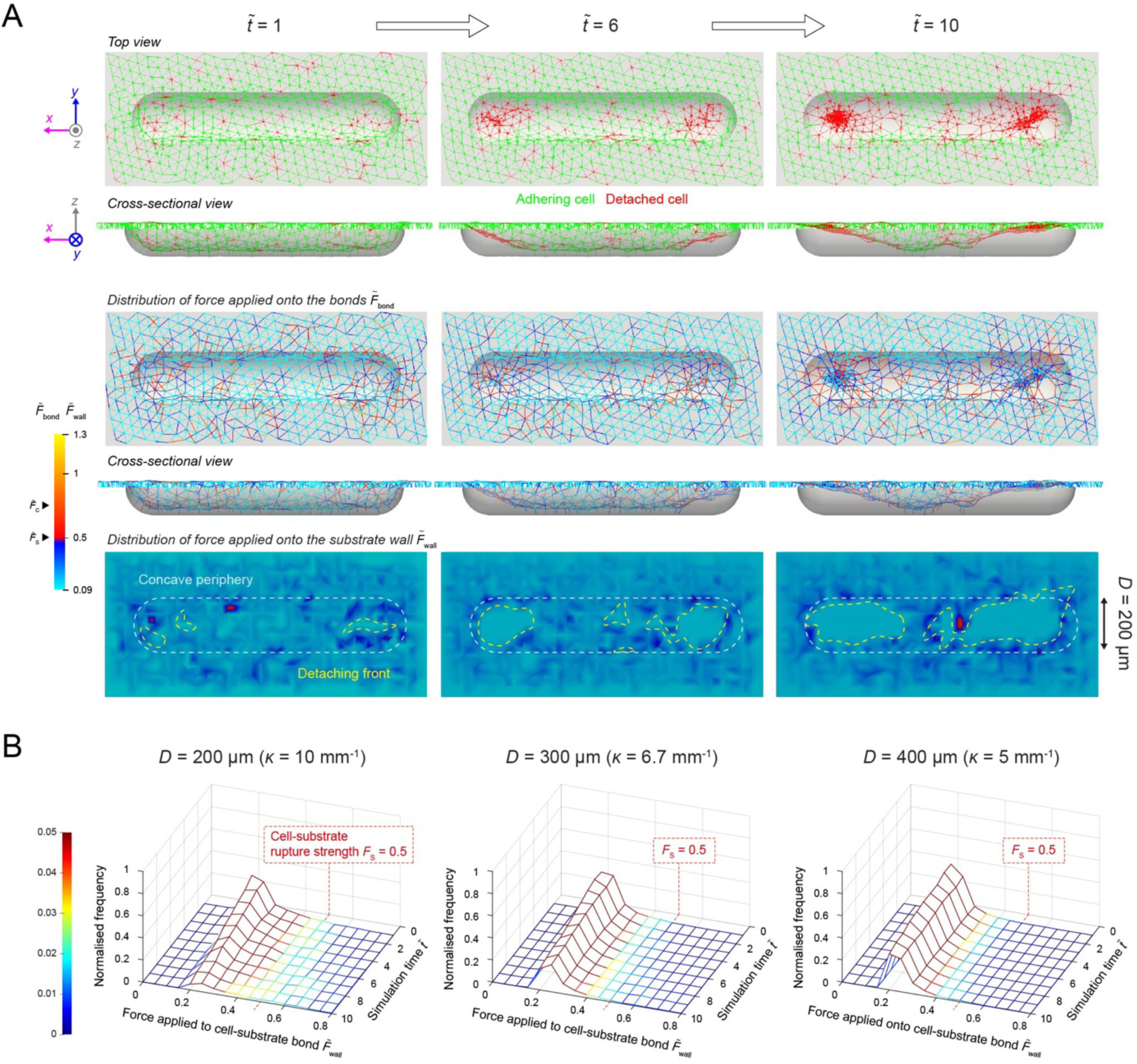
Cellular forces applied to the substrate surface *F*_wall_ simulated by the computational model. (A) Transition of cellular morphologies, and the spatial distribution of *F*_bond_ and *F*_wall_ throughout a microtissue detachment on a concavity with 200 µm of diameter. (B) Comparison of the transition of the frequency of *F*_wall_ applied to the concavities with different diameters over time (*D* = 200, 300 and 400 µm). Frequency of *F*_wall_ at each simulation time was normalised by the total number of cells located inside the concavity. The charts show the means of repeated simulations (*N* = 5).

To understand the details of detachment process, we next analysed the development of the range of *F*_wall_ occurrence frequencies over time, normalised by the total cell number inside the concavity (Fig. 7B). At the beginning of the simulation, the normalised frequency of *F*_wall_ showed a similar chart, having a lone peak at *F*_wall_ = 0.25 on all the concavities. But it should be highlighted that higher forces were present more often as the concavity diameter decreased from 400 to 200 µm, and the frequency of *F*_wall_ exceeding the rupture strength (*F*_S_ = 0.5) slightly increased from zero to a few percent. At the highest curvature (*D* = 200 µm), the chart rapidly flattened out over time; the peak height quickly dropped and the frequency of *F*_wall_ larger than *F*_S_ steadily increased. This indicates that more and more cellular contractile force was concentrated on the remaining cell-substrate bonds as the detachment progressed. It was also remarkable that the frequency of *F*_wall_ exceeding *F*_S_ remained less than 10% even at the peak of the detachment. In contrast, *F*_wall_ did not show a dramatic increase with time at lower curvatures (*D* = 300 and 400 µm), correlating to the suppressed detachment.

These results demonstrate that the concentration of cellular force at the cell-substrate interface plays a key role in the collective cellular detachment that essentially includes two distinct processes. First, surface curvature destabilises the cell-substrate adhesion, because the net cellular contractile force is applied in a direction normal to the cell-substrate interface. At the same time, the spatial variance in cellular density locally generates a fluctuation in the detaching force, which causes a stochastic rupture of cell-substrate adhesion. Second, the “defect” of cell adhesion is rapidly enlarged by the feedback loop of the successive concentration of the cellular contractile force on the detaching front, the break in the cell-substrate adhesion, and the shape optimisation of the detached cells; this acceleration process characterises the catastrophic nature of the detachment event. Substrate curvature boosts both the stochastic and catastrophic processes, accelerating the concentration of cellular contractility at the cell-substrate adhesion interface. It thus acts as an amplifier in the detachment event. The numerical analysis advocates an idea that the concentration of cellular contractile force needs to be carefully controlled by tuning the local curvature of the substrate and surface coatings when designing an *in vitro* environment to produce a microtissue with desirable morphology in three dimensions.

### 3.6. Particle-based model reproduces detachment behaviour of microtissues under complex three-dimensional constraints

Catastrophic microtissue detachment from tissue engineering scaffolds could be avoided with the help of quantitative prediction in tissue engineering. There is, however, no practical method of predicting the detachment behaviour of a microtissue from structured scaffolds. While smoothly curved substrates are suitable for examining geometry-guided cellular behaviours, micro-engineered scaffolds for cell culture often have an angulated shape due to limitations in conventional microfabrication techniques including soft lithography. We thus tested the applicability of our model to the practical prediction of microtissue morphology formed on such structures without curved planes. Using the proposed model, the detachment behaviour of microtissues from complex surfaces was simulated, and the results were compared with cell culture experiments.

Silicone elastomer substrates containing twelve-cornered-star-shaped holes were fabricated (Fig. 8A). These holes consisted of a flat bottom and vertical side walls without any curved planes and edges. Fibronectin was coated onto the substrate by either physisorption (PDL-fibronectin) or chemical crosslinking (SS-fibronectin) to promote initial cell adhesion. SMCs seeded onto the substrate formed a confluent sheet covering it, which subsequently ruptured and partially detached from the bottom surface. The detached microtissue typically formed a floating bridge connecting the centre part of the hole’s bottom and the top edge of the star-shaped cliffs (Fig. 8B). It was evident that the detachment frequently occurred at the corners of the holes. Overall, the structure of the bridging SMCs closely resembled those we categorised as “partial detachment” in the experiment using the concavity arrays (Fig. 3A).

**Fig. 8.**
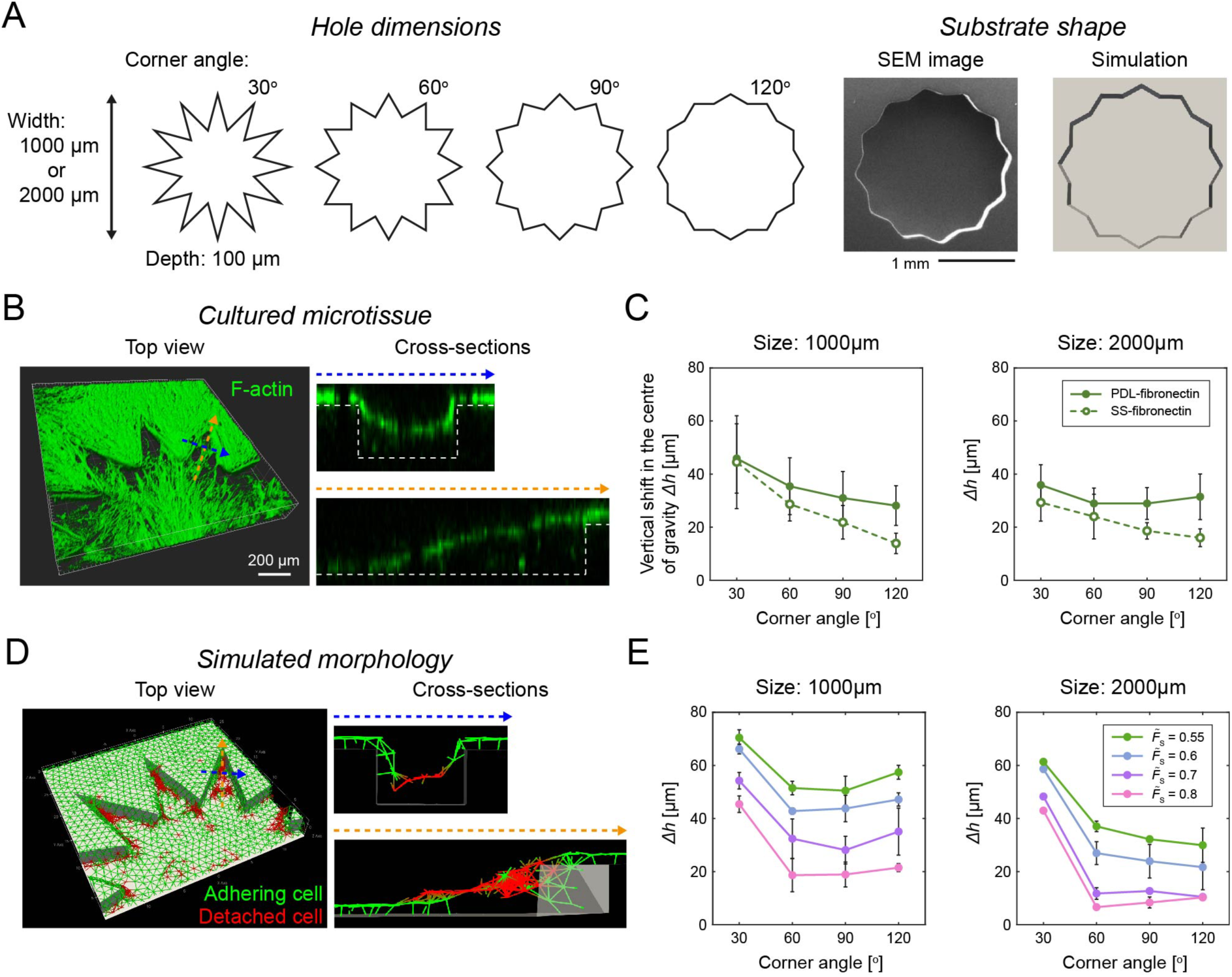
Comparison of the microtissue detached from star-shaped holes with the simulation result. (A) Dimension of the star-shaped holes. The structure was precisely fabricated on the PDMS surface. It was implemented in the simulation as a boundary condition. (B) Representative morphology of SMCs cultured on the star-shaped hole. (C) Analysis of microtissue detachment from the star-shaped holes with different corner angles (mean+SD, *N* = 3). Fibronectin was coated via either physisorption using PDL (PDL-fibronectin) or chemical crosslink using SulfoSANPAH (SS-fibronectin). (D) Microtissue morphology predicted by the simulation. (E) Predicted relationship of microtissue detachment and the corner angle at *ṫ* = 10. The rupture strength of cell-substrate bond *F*_S_ was varied (mean+SD, *N* = 3).

Focusing on each triangular region of the stars, the vertical shift in the centre of gravity *Δh* = *h* - *h*_A_ was quantified as an indicator of detachment. Larger *Δh* was observed as the corner angle decreased (Fig. 8C). This tendency qualitatively agrees with the observation that the higher substrate curvature induced more microtissue detachment from smoothly curved surfaces (Fig. 4), in the sense that a closer spatial confinement reduces the stability of the microtissue. Moreover, chemical crosslinking of fibronectin onto the substrate reduced *Δh*, indicating that the microtissue detachment was suppressed, similarly to what was observed on concavity arrays (Fig. 4A). The simulation reproduced the detachment process of a confluent SMC sheet, resulting in an identical morphology: SMCs hanging between the edges and the bottom of the star-shaped hole (Fig. 8D). It also predicted that a smaller corner angle promotes the progress of detachment, and that the effect is more evident in smaller holes (Fig. 8E). Our particle-based model provides a flexibility in defining substrate geometry not only for open substrates, but also for closed 3D structures. Based on an experimental study reporting a remodelling of epithelial cells in hollow hydrogel tubes [47], a bending tube (270 µm in tube inner diameter and 675 µm in curvature radius of the bending tube axis) that has both concave and convex within a unified geometry was set as a boundary condition. The simulation successfully reproduced the process in which cells detach from the outer side of the tube wall, shrink, and finally form a string on the inner surface of the tube (Fig. S5 and Supplementary movie 4), an identical dynamic reported in the previous study. The overall tendency predicted by the simulation agreed well with the experiments, yet there is room for fine tuning to fill the quantitative gap. This agreement supports the idea that our particle-based modelling approach is not limited in describing multicellular motion on ideally curved surfaces, but may have a much wider potential for predicting microtissue deformation on a wide range of structured scaffolds.

## 4. Discussion

Motivated by the rapid progress of biofabrication technologies, emerging microtissue-growth models have attracted a growing interest in rational scaffold design for tissue engineering [33,38,59]. The conventional models mainly describe the impact of surface curvature on the growth speed of confluent cell layers [32,33]. However, several experimental studies have demonstrated that the substrate curvature has another critical influence on cells; cellular contraction on a curved surface can induce spontaneous microtissue detachment [45–47]. Such cellular behaviour undermines the very basis of the tissue-growth models, but the detailed mechanism involved was poorly understood. This study, for the first time, clarifies the mechanism by which cellular contractile forces and geometrical constraints drive cellular detachment from structured surfaces, by combining experimental and computational approaches. The appearance probability maps of the morphologies of SMCs on trough-shaped concavities show that microtissue detachment from the scaffold surface is a highly stochastic process (Fig. 3). Specifically, the cellular detachment was promoted by large substrate curvature, enhanced cellular contractility, and weak cell-substrate adhesion strength, while it tended to be slightly suppressed by weak cell-cell adhesion (Fig. 4), confirming the basic tendency reported in the previous experimental studies [45–47]. Established disciplines applied for explaining unique cellular behaviours on curved surfaces, such as continuum [32], phase-field [60], vertex [61], particle [62] and interface [33] had limited flexibility in depicting stochastic rupture, topological change, and large deformation of the microtissue body, the key features of microtissue detachment. We thus took a new modelling approach, translating cells into discrete particles. Our particle-based model is unique in explicitly describing the stochastic adhesion and rupture between cells as well as those between a cell and the substrate. This particle-based approach allowed us to naturally describe topological change and large deformation of the microtissue body. Also, the key parameters, the amplitude modulus of cell contraction *α*, the rupture strength of cell-cell bond *F*_C_, and that of cell-substrate bond *F*_S_ are explicit and intuitive enough to compare with experimental conditions. Despite its simplicity, the model enabled us to reproduce the typical morphological changes undergone by SMCs on curved surfaces (Fig. 5) as well as the basic dependency of the cellular detachment to substrate curvature, cell-substrate adhesion strength and cellular contractility (Fig. 6). The simulations quantitatively elucidated the mechanics of how stress concentration at the cell-substrate interface drives stochastic and catastrophic microtissue detachment (Fig. 7). By virtue of its flexibility in defining a complex substrate geometry as a boundary condition, the model successfully predicted how a microtissue detaches inside star-shaped holes without curvature (Fig. 8). This is, to the best of our knowledge, the first attempt to describe the complex microtissue dynamics including rupture and detachment on structured surfaces on a physical basis.

Highlighting relaxation-based morphological changes in microtissues, our computational model reproduced the basic detachment and aggregation processes of SMCs on structured surfaces to a certain extent. It should nevertheless also be mentioned that there is still a slight gap between the simulations and the observations, both qualitatively and quantitatively. Qualitatively, the simulated cellular motion is in principle passive; experimental observations showed that a group of cells detached partially and ruptured simultaneously with migration, followed by aggregation in the end (Fig. 3A and Supplementary movie 1), whereas simulations predicted that cells first detached from the substrate in a sheet form, ruptured, and then aggregated (Fig. 5A and supplementary movie 2). Quantitatively, there is still a modest difference of around 1.5 times between the experimental and the predicted values in *Δh* (Fig. 8 C and E). We think two major causes underly these differences. First, the current model does not consider some microscopic features of SMCs such as highly stretched cellular shape and local cellular migration. Second, our model only considers the microtissue shape in force equilibrium and underestimates the viscous nature of cell bodies during rupture and deformation. Also, the difference in the time scale of cell-cell and cell-substrate adhesions is not considered. The individual contributions of these different types of adhesion, which are likely to influence the effective viscosity of the cell layer, could be distinguished by introducing different time-correction terms for each into the bond-rearrangement functions expressed by Eqs. 9 and 10. Consideration of viscous resistance as well as accurate representations of microscopic cellular events such as adhesion, separation and migratory motion might be the key to a precise description of the transient microtissue morphology from a sheet to a clump in three dimensions.

In the current study, we experimentally revealed the influence of cellular contractility, rupture strength of cell-substrate and cell-cell adhesions, and substrate curvature onto the morphology of SMCs by using drugs, a chemical crosslinker, antibody, and trough-shaped concavities with different scales (Fig. 3 and 4). It should be noted that these parameters tuned in the experiments potentially interact with each other and cannot be tuned independently. For example, drug treatments affecting the cellular contractility may have altered the rupture strength of cell-substrate adhesion because of the mutual interaction between cellular force and mechanosensitive molecular complex, *e.g.*, focal adhesions [63], formed at the cell-substrate interface. Also, substrate curvature at this supracellular scale is recently known to influence cell differentiation [64,65], which potentially changes the magnitude of cell contraction. Such interdependency of the experimental parameters may have resulted in some minor features of the probability maps that our model did not predict; there is a constant probability of adhesion over the different curvatures in the presence of TGF-β (Fig. 3D) and SMCs showed detachment at lower curvature with a few percent of probability even in the presence of blebbistatin (Fig. 3E). In addition, experimental characterisation of the influence of cell-cell adhesion onto the detachment event remains a challenge in future investigations. Increase in antibody concentration up to 100 µg/mL scale [66] or knockdown of N-cadherin expression might contribute to the quantitative assessment of the role of cell-cell interactions. While the basic tendency of cell sheet detachment has been explained on a physical basis, the impact of hidden changes in biological parameters remains to be elucidated based upon a precise tuning of biophysical parameters, to fully understand the details of microtissue deformation.

Nevertheless, the overall consistency between the simulation and experiments on microtissue detachment supports the idea that the macroscopic balance of substrate curvature, cellular contraction and cell-substrate adhesion strength at the cell-substrate interface is a key to predicting whether cells will succeed in forming a stable layer on structured scaffolds. Bakarat *et al.* reported that contractile forces mutually applied onto SMCs within a confluent layer generate a hole in the cell sheet and this defect initiates the subsequent collective clustering and condensation of cells on a flat surface [67]. Our model suggests that 3D structures of the substrate, including curvature, promote such contractility-driven events by amplifying the stress concentration at the cell-substrate adhesion interface, underlining the increasing importance of surface treatment in 3D scaffold design. Brochard-Wyart *et al.* explained cellular aggregation and spread based on the minimisation of interfacial energy, *i.e.*, surface affinity [55]. Our model, describing microtissue shape by finding the local minimum of the effective energy of cellular contraction and size exclusion, is compatible with their notion. Thus, cells in our simulation naturally form a clump when *F*_C_ is larger than *F*_S_ (Fig. 5), in good accordance with their affinity-based description. On the other hand, our model explicitly considers force-dependent rupture and geometry-dependent adhesion processes during the microtissue deformation. These stochastic operations allowed us to describe irreversible and route-dependent microtissue deformation processes, which an affinity-based framework has fundamental difficulty reproducing. Together with the flexibility in defining the substrate geometry, our model could be positioned as an engineering-oriented extension of the conventional soft-matter-based microtissue model.

From an engineering perspective, our study helps bioengineers design micro-engineered scaffolds to control microtissue morphology. In the field of cell sheet engineering, it has been qualitatively known that spontaneous cell-sheet detachment from a thermoresponsive polymer surface requires cell-generated force and cytoskeletal reorganisation [68]. But the mechanical principle has been poorly understood for a long time. In the present study, we clarified that the progressive stress concentration critically impairs the cell-substrate adhesion interface. Quantitative knowledge is particularly helpful when designing scaffolds that actively utilise cellular detachment to engineer microtissues in three dimensions. For example, collective cellular detachment can be, if controlled, employed to create microtissues in unique 3D morphologies, such as strings [69] and bridges between piers [70]. Our modelling framework is in principle applicable to simulate cellular detachment from a scaffold in confined 3D environments, *e.g.*, tubular structures (Fig. S5 and Supplementary movie 4), and not limited to open substrates. But fine tuning of parameters as well as implementation of cell-type-specific microscopic features including anisotropic cellular geometry and size distribution is required for precise description, because subtle changes in the related parameters would critically influence the stress concentration in complex 3D environments. These are the future challenges to predict accurate microtissue deformation processes on fully 3D scaffolds with complex geometry that the currently proliferating 3D bioprinting techniques provide for tissue engineering. The mechanical understanding shown in this study would give deep insights into various aspects of scaffold design in tissue engineering, including scaffold geometry, surface treatments and growth factors. Also, the proposed computational framework provides a key step towards microtissue-growth models being able to predict whether cells will form a stable monolayer on structured surfaces in the initial stage of the microtissue formation process. These insights in microtissue detachment and deformation on structured surfaces, as revealed by our combined approach, would greatly expand the potential applicability of the emerging microtissue modelling frameworks, and thus might contribute to an effective and rational scaffold design for tissue engineering in the future.

## 5. Conclusion

This study, for the first time, clarified the mechanism underlying spontaneous microtissue detachment from structured surfaces. The series of observations of SMCs cultured on trough-shaped concavities revealed how substrate curvature, cell-substrate adhesion strength and cellular contractility influence the 3D morphologies of SMCs. A new particle-based model that explicitly describes how cellular contractile forces cause rupture and deformation of microtissue was proposed. The model-based simulations reproduced the morphologies and deformation process of SMCs as well as their dependencies on mechanical parameters; the correspondence with the experiments clarified that a stress concentration at the cell-substrate interface drives stochastic and catastrophic microtissue detachment. This method would enable tissue engineers to predict microtissue dynamics on complex microstructures on a physical basis. These understandings as well as our unique modelling framework provide a new platform for designing scaffolds that properly guide the engineered tissue formation process. This study thus contributes to a rational scaffold design in tissue engineering and regenerative medicine in the future.

## Acknowledgements

The authors acknowledge funding from Japan Society for the Promotion of Science (JSPS), KAKENHI Grant Numbers (JP18H5963, JP19K20679,21K18041, 20K20958 and 21H01209) and Bilateral Program Grant Number JPJSBP1201976; Japan Science and Technology Agency (JST), CREST Grant Number JPMJCR1921; Japan Agency for Medical Research and Development (AMED), Grant Number 21bm0704065h0001; The World Premier International Research Center Initiative, Ministry of Education, Culture, Sports, Science and Technology (MEXT), Japan; Mizuho Foundation for the Promotion of Sciences.

## Supplementary data

Supplementary figures S1, S2, S3 and S4 and table S1 are provided in a separate document.

Supplementary movies S1, S2, S3 and S4 are attached as separate video files.

## Notes

### Competing Interest Statement

The authors have declared no competing interest.

### Summary of Updates

Title revised; Authors updated; Experiments and simulations testing the influence of cell-cell adhesion strength added; Figure 3, 4, 6 and 7 revised; Supplementary files updated;

