## Supplementary information for "Multicellular dynamics on structured surfaces: Stress concentration is a key to controlling complex microtissue morphology on engineered scaffolds"

**Supplementary figures**

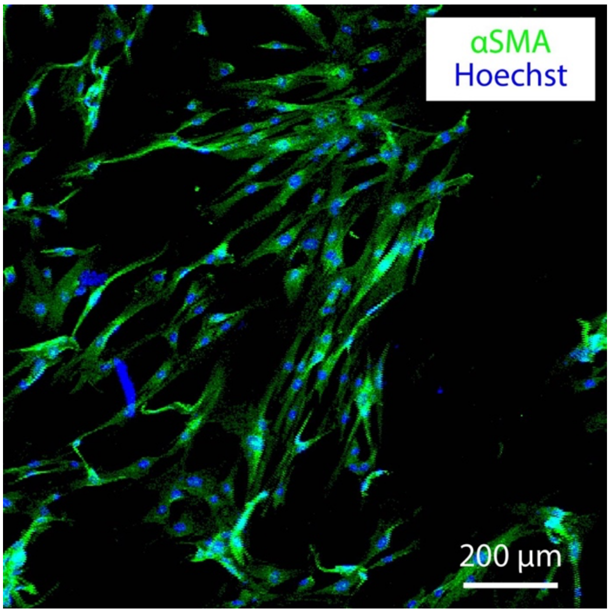


Fig. S1. Observation of αSMA, contractile phenotype marker of SMC.

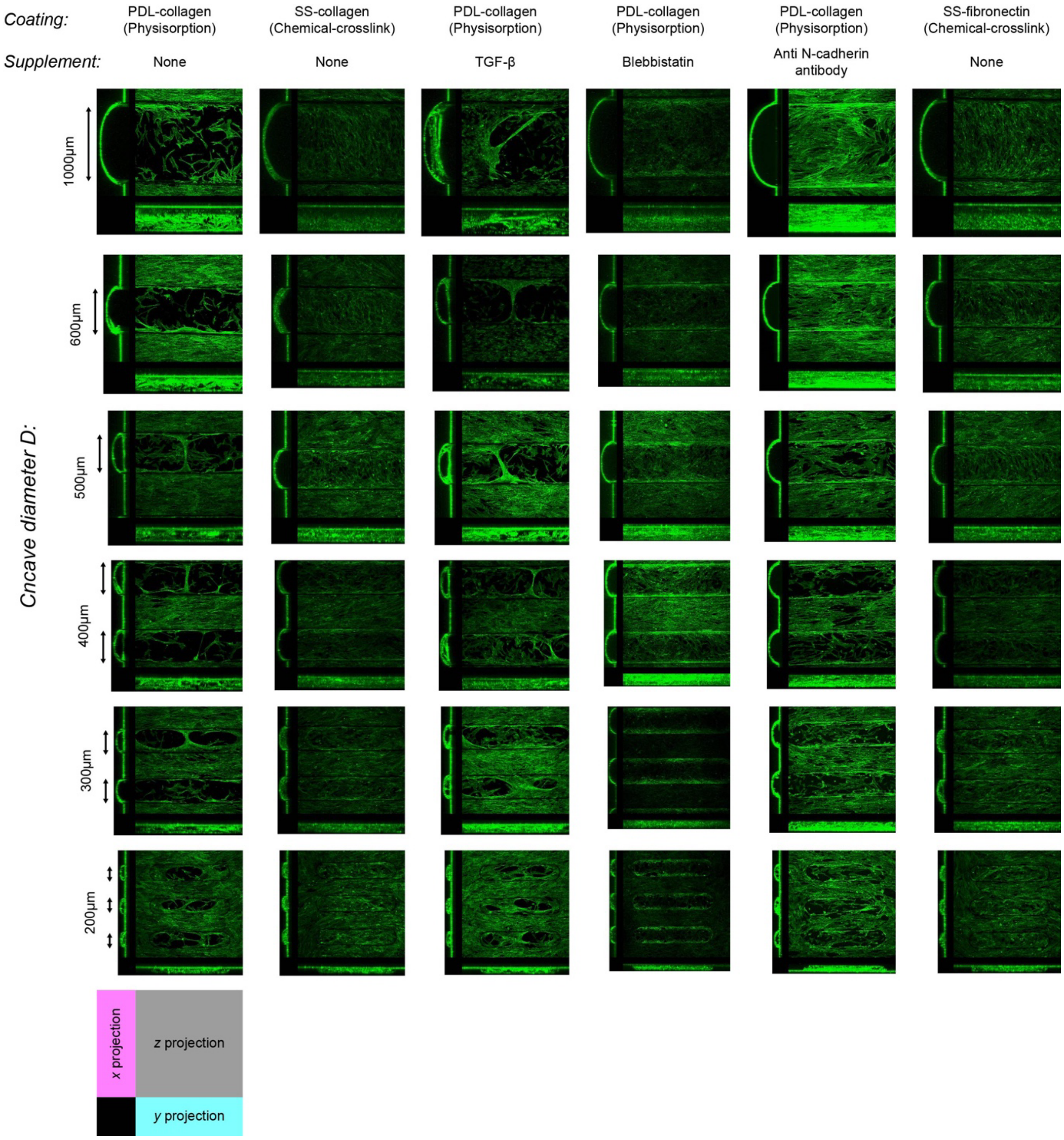


Fig. S2. Overview of the morphologies of SMCs cultured on trough-shaped concavity arrays with different substrate coatings and supplements. F-actin was stained and observed via confocal microscopy.

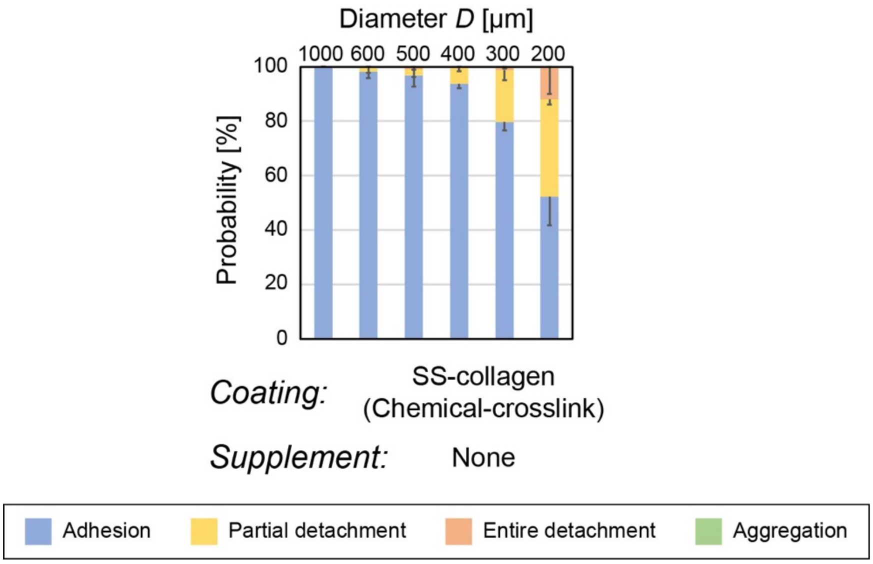


Fig. S3. Appearance probability map of the four representative morphologies of SMCs cultured on concavities coated with fibronectin via chemical crosslink (mean-SE, $N = 3$).

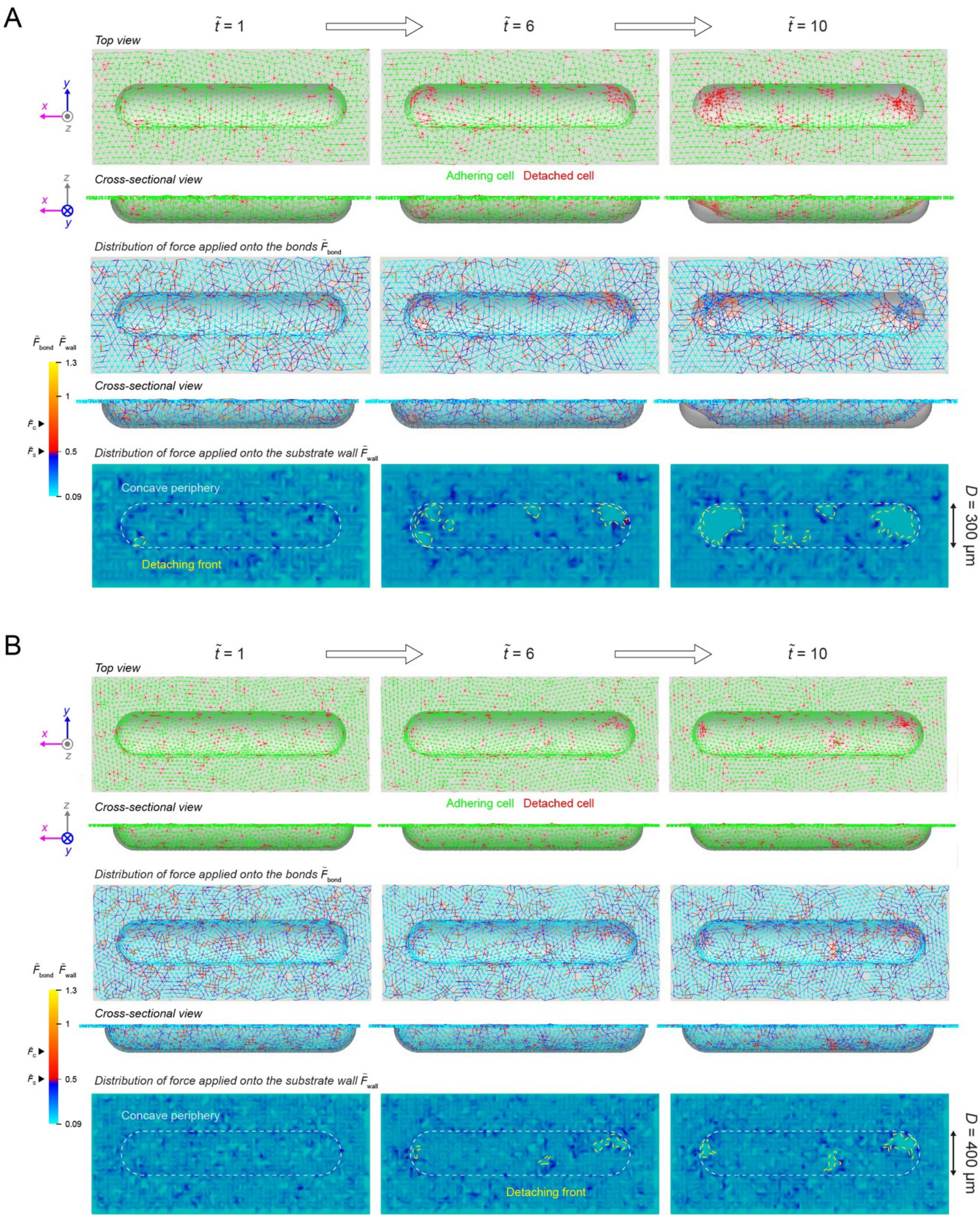


Fig. S4. Simulated cellular morphologies, and spatial distribution of $F_{\mathrm{bond}}$ and $F_{\mathrm{wall}}$ on concavities with lower curvatures. (A) $D$ = 300 μm ($\kappa$ = 6.7 mm$^{-1}$). (B) $D$ = 400 μm ($\kappa$ = 5 mm$^{-1}$).

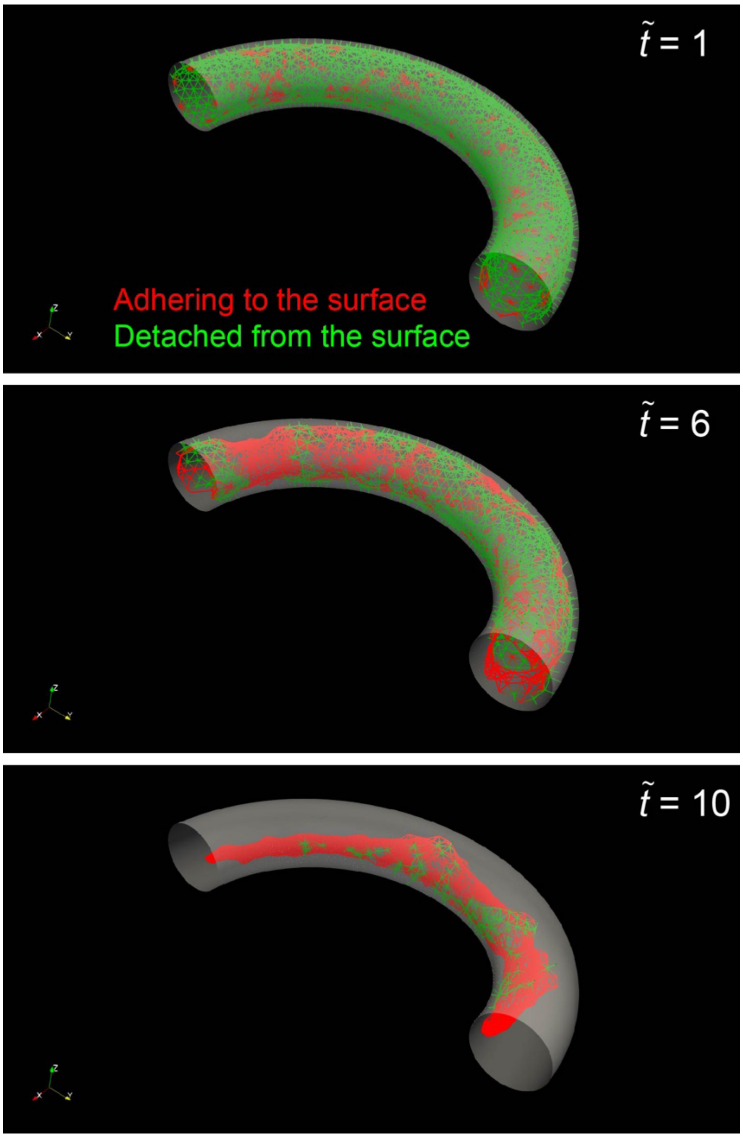


Fig. S5. Simulated cellular detachment from the inner surface of a bending tubular structure (tube diameter: 270 µm, curvature radius of bending tubular axis: 675 µm).

### Supplementary table

Supplementary table S1. List of the values assigned to the simulation parameters

| Figure | $\alpha$ | $\tilde{F}_S$ | $\tilde{F}_C$ | Substrate geometry |
|---|---|---|---|---|
| Fig. 5 (Supplementary movie 2) | 0.8 | 0.5 | 0.75 | Trough-shaped concavity<br>$D$ = 200 µm |
| Fig. 6A, 7, S4 (Supplementary movie 3) | 0.8 | 0.5 | 0.75 | Trough-shaped concavity<br>$D$ = 200, 300, 400 µm |
| Fig. 6B | 0.8 | Varied from 0.45 to 0.6 | 0.75 | Trough-shaped concavity<br>$D$ was varied from 200 to 1000 µm |
| Fig. 6C | Varied from 0.7 to 0.9 | 0.5 | 0.75 | Trough-shaped concavity<br>$D$ was varied from 200 to 1000 µm |
| Fig. 6D | 0.8 | 0.5 | Varied from 0.6 to 0.9 | Trough-shaped concavity<br>$D$ was varied from 200 to 1000 µm |
| Fig. 6E | Varied from 0.7 to 0.9 | Varied from 0.45 to 0.6 | 0.75 | Trough-shaped concavity<br>$D$ = 200, 300, 400 µm |
| Fig. 6F | 0.8 | Varied from 0.45 to 0.6 | Varied from 0.6 to 0.9 | Trough-shaped concavity<br>$D$ = 200, 300, 400 µm |
| Fig. 8D | 0.8 | 0.8 | 0.75 | Star-shaped hole<br>30° corner angle<br>2000 µm width |
| Fig. 8E | 0.8 | Varied from 0.55 to 0.8 | 0.75 | Star-shaped hole<br>The corner angle was varied from 30 to 120°<br>The width was 1000 or 2000 µm |
| Fig. S5 (Supplementary movie 4) | 0.8 | 0.5 | 0.75 | Bending tube<br>Tube inner diameter was 270 µm<br>The curvature radius of the bending tube axis was 675 µm |

**Supplementary movies**

Supplementary movie 1. Deformation of SMCs on the concavity:

Time-lapse movie of SMCs that spontaneously detach and aggregate on a trough-shaped concavity ($D$ = 1000 μm).

Supplementary movie 2. Simulated microtissue deformation:

Simulated microtissue deformation on a trough-shaped concavity ($D$ = 200 μm).

Supplementary movie 3. Simulated cellular traction force applied to the concavity surface:

Simulated microtissue deformation on a trough-shaped concavity ($D$ = 200 μm) and the spatial distribution of cellular traction force applied to the concavity surface $F_{\text{wall}}$.

Supplementary movie 4. Simulated cellular detachment inside a bending tube:

Simulated cellular detachment from the inner surface of a bending tubular structure (tube inner diameter: 270 μm, curvature radius of bending tubular axis: 675 μm).
